# PAX3-FOXO1 drives targetable cell state-dependent metabolic vulnerabilities in rhabdomyosarcoma

**DOI:** 10.1101/2025.01.15.633227

**Authors:** Katrina I. Paras, Julia S. Brunner, Jacob A. Boyer, Angela M. Montero, Benjamin T. Jackson, Sangita Chakraborty, Abigail Xie, Kristina Guillan, Armaan Siddiquee, Lourdes Pajuelo Torres, Joshua D. Rabinowitz, Andrew Kung, Daoqi You, Filemon Dela Cruz, Lydia W. S. Finley

## Abstract

PAX3-FOXO1, an oncogenic transcription factor, drives a particularly aggressive subtype of rhabdomyosarcoma (RMS) by enforcing gene expression programs that support malignant cell states. Here we show that PAX3-FOXO1^+^ RMS cells exhibit altered pyrimidine metabolism and increased dependence on enzymes involved in *de novo* pyrimidine synthesis, including dihydrofolate reductase (DHFR). Consequently, PAX3-FOXO1^+^ cells display increased sensitivity to inhibition of DHFR by the chemotherapeutic drug methotrexate, and this dependence is rescued by provision of pyrimidine nucleotides. Methotrexate treatment mimics the metabolic and transcriptional impact of *PAX3-FOXO1* silencing, reducing expression of genes related to PAX3-FOXO1-driven malignant cell states. Accordingly, methotrexate treatment slows growth of multiple PAX3-FOXO1^+^ tumor xenograft models, but not fusion-negative counterparts. Taken together, these data demonstrate that PAX3-FOXO1 induces cell states characterized by altered pyrimidine dependence and nominate methotrexate as an addition to the current therapeutic arsenal for treatment of these malignant pediatric tumors.

## Introduction

Cellular metabolic pathways support critical cell functions, ranging from macromolecule biosynthesis and energy production to maintenance of redox homeostasis and cell signaling pathways. As cellular functions change, so does the demand for reducing equivalents, energy production, and macromolecule synthesis. Accordingly, cellular metabolic demands are often highly context-dependent, with cells of different lineages or even in distinct stages of differentiation experiencing unique metabolic phenotypes^1–3^. Consequently, metabolic perturbations that are fatal to one cell state may be well-tolerated in another. This context-specific role of metabolic programs is exemplified by inborn errors of metabolism, a constellation of human disorders driven by germline mutation in metabolic enzymes in which disease severity and affected organ system vary depending on the metabolic pathway—and even, the specific step within a metabolic pathway—that is mutated^4,5^. The notably heterogeneous clinical manifestations of inborn errors in metabolism underscores both how different cell states respond differently to metabolic alterations and how modifying factors including genotype and diet can affect response to metabolic perturbations^5^.

The observation that many cell states have unique metabolic requirements raises the possibility that these metabolic vulnerabilities can be targeted to eliminate specific cell types. In cancer, such metabolic targeting has notable successes^6^. For example, pediatric acute lymphoblastic leukemia is markedly sensitive to depletion of asparagine through asparaginase therapy^7^, and antifolates such as aminopterin and methotrexate are classic agents for acute lymphoblastic leukemia^8^, choriocarcinoma^9^, and osteosarcoma^10^. To date, many metabolic therapies aim to target the rapid proliferation of cancer cells or to exploit collateral lethality secondary to genetic mutations harbored by tumor cells^11,12^. Whether different cell states that emerge within and between tumor subtypes harbor different metabolic vulnerabilities is relatively poorly understood.

Here, we aimed to determine whether metabolic dependencies could be leveraged to target malignant cell states using rhabdomyosarcoma (RMS) as a model system. RMS is the most common soft tissue sarcoma in children and shares histological and transcriptional signatures of normal myogenic development^13^. RMS largely falls into two subtypes based on the presence or absence of a fusion oncogene in which the DNA-binding domain of *PAX3* or, more rarely, *PAX7*, is fused with the transactivation domain of *FOXO1*^14^. Whereas fusion-negative tumors are more likely to be low-risk and thus effectively managed with a combination of chemotherapy, surgery, and radiotherapy^15^, fusion-positive tumors—especially those expressing the PAX3-FOXO1 fusion—are higher risk and more likely to be metastatic at diagnosis^16,17^. Accordingly, the presence of PAX3-FOXO1 correlates with worse outcomes for patients with either localized or metastatic disease, where 5-year overall survival is as low as 13% compared to 62% for fusion-negative disease^18,19^. Despite these different prognoses, the therapeutic approach to fusion-positive tumors is identical to fusion-negative tumors and has remained largely unchanged for over 30 years^13^.

Recent single-cell profiling of RMS cell lines and tumors revealed that RMS cells adopt a spectrum of different cell states that may contribute to malignant progression. Specifically, fusion-negative RMS cells retain the ability to access a range of cell states across the normal myogenic developmental cascade; in contrast, fusion-positive RMS cells are locked into a “cycling progenitor” state that is resistant to normal myogenic differentiation and associated with poor prognosis^20,21^. By comparing metabolic dependencies of fusion-positive and fusion-negative RMS cell lines, we aimed to determine whether the malignant cell states that characterize PAX3-FOXO1^+^ RMS have metabolic vulnerabilities that can be exploited therapeutically for this hard-to-treat disease.

## Results

### Rhabdomyosarcoma subtypes harbor intrinsic metabolic differences

We first aimed to determine whether cultured RMS cell lines recapitulate cell state differences observed in human tumors. RMS cell lines profiled by the DepMap project^22^ were grouped by fusion status and analyzed for expression of genes characterizing the muscle stem cell (MuSC) or differentiated populations typifying fusion-negative tumors and the cycling progenitor and cell cycle (S-phase) state that distinguish fusion-positive tumors^20^. Genes upregulated in PAX3-FOXO1^+^ relative to fusion-negative cell lines were significantly enriched for genes belonging to the cycling progenitor and cell cycle state; in contrast, genes upregulated in fusion-negative cell lines were enriched for MuSC and, to a lesser extent, differentiation-associated genes (Supplementary Fig. 1A). Thus, although cultured cell lines cannot fully capture the phenotypic heterogeneity present in human tumors, RMS cell lines do recapitulate aspects of the aberrant cell fate programs observed in PAX3-FOXO1^+^ relative to fusion-negative RMS tumors.

Next, we asked whether metabolic profiles varied with RMS subtype using a panel of two PAX3-FOXO1^+^ RMS lines (RH30, RH41) and two fusion-negative RMS lines (RD, SMS-CTR). Unsupervised analysis of liquid chromatography-mass spectrometry (LC-MS)-based metabolomic profiling of 126 metabolites revealed that cell lines clustered based on fusion status (Fig. 1A). Of the 126 metabolites analyzed, 66 demonstrated significant changes (*P* < 0.05) between subtypes (Supplementary Fig. 1B). To identify metabolites that reliably distinguish PAX3-FOXO1^+^ RMS cells from fusion-negative RMS cells, we performed supervised clustering using variable importance in the projection (VIP) analysis (Supplementary Fig. 1C). Overall, metabolomic changes were largely reproducible, with 29 metabolites emerging with VIP scores > 1.0 in two independent LC-MS experiments (Fig. 1B). Of these metabolites, intermediates in pyrimidine metabolism and pyrimidine nucleotides were consistently elevated in both PAX3-FOXO1^+^ RMS cell lines compared to their fusion-negative counterparts (Fig. 1C, D).

**Figure 1.**
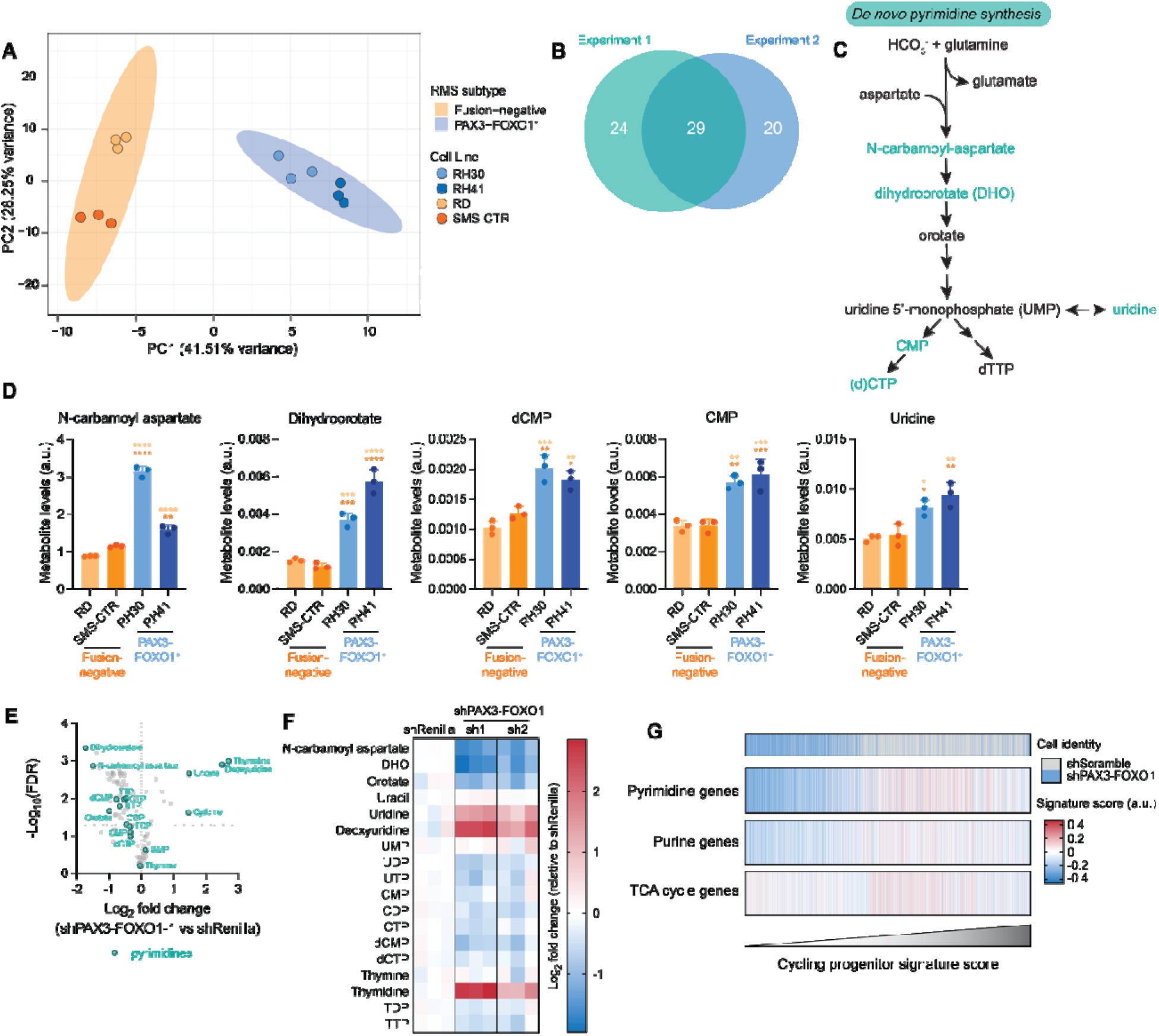
Rhabdomyosarcoma subtypes harbor intrinsic metabolic differences. **A**, Principal component analysis of targeted metabolomics data of 126 metabolites obtained from four RMS cell lines. Ellipses represent 95% confidence intervals within each sample group. **B**, Venn diagram depicting metabolites that discriminate between PAX3-FOXO1^+^ and fusion-negative RMS across two independent LC-MS experiments. Metabolites with variable importance in the projection (VIP) scores over 1.0 were included in the analysis. **C**, Schematic depicting *de novo* pyrimidine synthesis pathway. **D**, Levels of N-carbamoyl aspartate, dihydroorotate (DHO), deoxycytidine monophosphate (dCMP), cytidine monophosphate (CMP), and uridine. **E**, Volcano plot showing log_2_ fold change in metabolite abundance in RH30 cells expressing shRNA against *PAX3-FOXO1* compared to cells expressing shRNA against *Renilla* luciferase (control). Cells were cultured with doxycycline for 72 h to induce shRNA expression. Pyrimidine metabolites are highlighted in teal. Dotted horizontal line indicates significance threshold FDR < 0.05. **F**, Heatmap depicting levels of pyrimidine metabolites in RH30 cells expressing shRNAs against *PAX3-FOXO1*, expressed as the log_2_-transformed fold change relative to shRenilla. Data are from same experiment as data shown in **E**. **G**, Heatmap of gene signature scores for pyrimidine, purine, and TCA cycle genes obtained from published single-cell RNA-sequencing data^20^ of KFR RMS cells expressing shRNA against *PAX3-FOXO1* (shScramble serves as a control). Cells were ranked in order of increasing cycling progenitor signature score. Gene lists are provided in Supplementary Table 3. For **D**, data are mean ± s.d., *n* = 3 independent replicates. For **D**, statistical significance was assessed by ordinary one-way ANOVA with Tukey’s multiple comparisons test, with each PAX3-FOXO1^+^ cell line compared to each fusion-negative cell line. (\**P* < 0.05, \*\**P* < 0.01, \*\*\**P* < 0.001, \*\*\*\**P* < 0.0001)

To determine whether the observed metabolic changes were a consequence of fusion status, we introduced doxycycline (dox)-inducible hairpins targeting *PAX3-FOXO1* into RH30 cells. Three days of dox administration was associated with robust loss of PAX3-FOXO1 and a modest decline in proliferation (Supplementary Fig. 1D, E), consistent with previous work demonstrating that only prolonged PAX3-FOXO1 repression induces growth arrest and/or cell death^23,24^. Metabolomic profiling revealed that PAX3-FOXO1 silencing triggered changes in many metabolites related to pyrimidine metabolism, with notable downregulation in intermediates in the *de novo* pyrimidine biosynthesis pathway (Fig. 1E, F). The observation that N-carbamoyl aspartate and dihydroorotate, intermediates in pyrimidine biosynthesis decreased in fusion-negative cells, are among the most downregulated metabolites following PAX3-FOXO1 silencing is consistent with the notion that pyrimidine biosynthesis is closely tied to PAX3-FOXO1 expression.

To determine whether changes in pyrimidine metabolism are a direct consequence of transactivation by PAX3-FOXO1 or secondary to changes in cell state driven by PAX3-FOXO1 expression, we interrogated published datasets delineating direct PAX3-FOXO1 targets. Across five available datasets profiling PAX3-FOXO1 chromatin occupancy by chromatin immunoprecipitation followed by sequencing (ChIP-seq)^25–27^, known PAX3-FOXO1-target genes were reproducibly identified as directly bound by PAX3-FOXO1 (Supplementary Fig. 1F). In contrast, no genes involved in pyrimidine synthesis emerged as reproducible targets of PAX3-FOXO1 (Supplementary Fig. 1F). We therefore examined the alternate hypothesis that pyrimidine metabolism varies alongside RMS cell state. Analysis of transcriptional profiling of RH30 cells following acute depletion of PAX3-FOXO1 (ref^24^) revealed that PAX3-FOXO1 controls RMS cell state in cultured cells: following PAX3-FOXO1 degradation, genes related to fusion-negative cell states (MuSC, differentiated) were largely upregulated while genes related to fusion-positive cell states (cycling progenitor, S-phase) were largely downregulated (Supplementary Fig. 1G, H). Of the genes gradually downregulated following PAX3-FOXO1 degradation were three genes crucial for *de novo* pyrimidine synthesis: the trifunctional enzyme carbamoyl-phosphate synthetase 2, aspartate transcarbamylase and dihydroorotase (*CAD*), thymidylate synthase (*TYMS*) and dihydrofolate reductase (*DHFR*) (Supplementary Fig. 1I). Both DHFR and TYMS were likewise downregulated following expression of shRNA targeting *PAX3-FOXO1* in two independent fusion-positive RMS cell lines, whereas CAD showed inconsistent effects across cell lines (Supplementary Fig. 1J). To more broadly investigate the expression of genes involved in metabolic pathways across different cell states in these datasets, we ranked cells assessed by single cell-RNA-sequencing^20^ by their cycling progenitor gene signature score, confirming that cells expressing a hairpin targeting PAX3-FOXO1 overwhelmingly exhibited low scores for this PAX3-FOXO1-associated cell state (Fig. 1G). Consistently, we found that cells exhibiting low cycling progenitor gene scores also exhibited lower levels of genes associated with pyrimidine synthesis, and to a lesser extent, purine synthesis, but not genes in unrelated metabolic pathways such as the tricarboxylic acid (TCA) cycle (Fig. 1G). Together, these results indicate that altered expression of pyrimidine related genes and altered pyrimidine metabolites are intrinsic features of PAX3-FOXO1^+^ cell states.

### Genome-wide CRISPR screens reveal metabolic vulnerabilities specific to PAX3-FOXO1^+^ RMS

The above findings raise the possibility that transcriptional heterogeneity between malignant cell states in RMS is accompanied by distinct metabolic programs. To determine whether the metabolic differences inherent to PAX3-FOXO1^+^ and fusion-negative RMS subtypes result in subtype-specific metabolic vulnerabilities, we leveraged genome-wide CRISPR screens generated through the DepMap project^22^. For every gene in the genome, the average essentiality score from 5 fusion-negative RMS lines was subtracted from the average essentiality score from 7 PAX3-FOXO1^+^ RMS lines, resulting in a differential essentiality score (Supplementary Table 1). Ranking genes by their differential essentiality revealed that *PAX3* is the most differentially essential gene between the two RMS subtypes (Fig. 2A), likely because the sgRNAs targeting *PAX3* also target the *PAX3-FOXO1* fusion. Further validating the approach, we found that *MYOG*, a muscle differentiation marker known to be more highly expressed in PAX3-FOXO1^+^ cells and tumors^28^, was the 2^nd^ most differentially essential gene. Among metabolic genes, *DHFR* and *TYMS* emerged as the 1^st^ and 3^rd^ most differentially essential, respectively (Fig. 2B). Even when compared to all genes in the genome, *DHFR* ranked 9^th^ and *TYMS* ranked 12^th^, ahead of *MYCN* (17^th^), an oncogenic driver with a well-established role in many cases of PAX3-FOXO1^+^ RMS^27,29^ (Fig. 2A). The differential dependence on DHFR/TYMS was not simply driven by the fusion-negative RMS lines being unusually resistant to DHFR/TYMS inhibition: compared to all lines in DepMap, PAX3-FOXO1^+^ RMS lines are more sensitive to loss of DHFR/TYMS, while fusion-negative RMS lines are not (Fig. 2C). Although DHFR and TYMS are involved in the folate cycle and pyrimidine biosynthesis, respectively, the two enzymes are tightly linked through dihydrofolate (DHF), which is both the product of TYMS enzymatic activity and the substrate for production of tetrahydrofolate (THF) by DHFR (Fig. 2D). DHF inhibits TYMS by negative feedback inhibition, and DHF accumulation is associated with cellular toxicity; accordingly, DHFR activity is required for sustained TYMS function^30^. Indeed, in many bacterial species, DHFR and TYMS form a tightly coupled enzymatic unit in which TYMS activity cannot exceed that of DHFR^31^. Accordingly, the emergence of DHFR and TYMS as top dependencies in PAX3-FOXO1^+^ supports the hypothesis that dependence on *de novo* thymidine synthesis defines malignant PAX3-FOXO1^+^ cell states.

**Figure 2.**
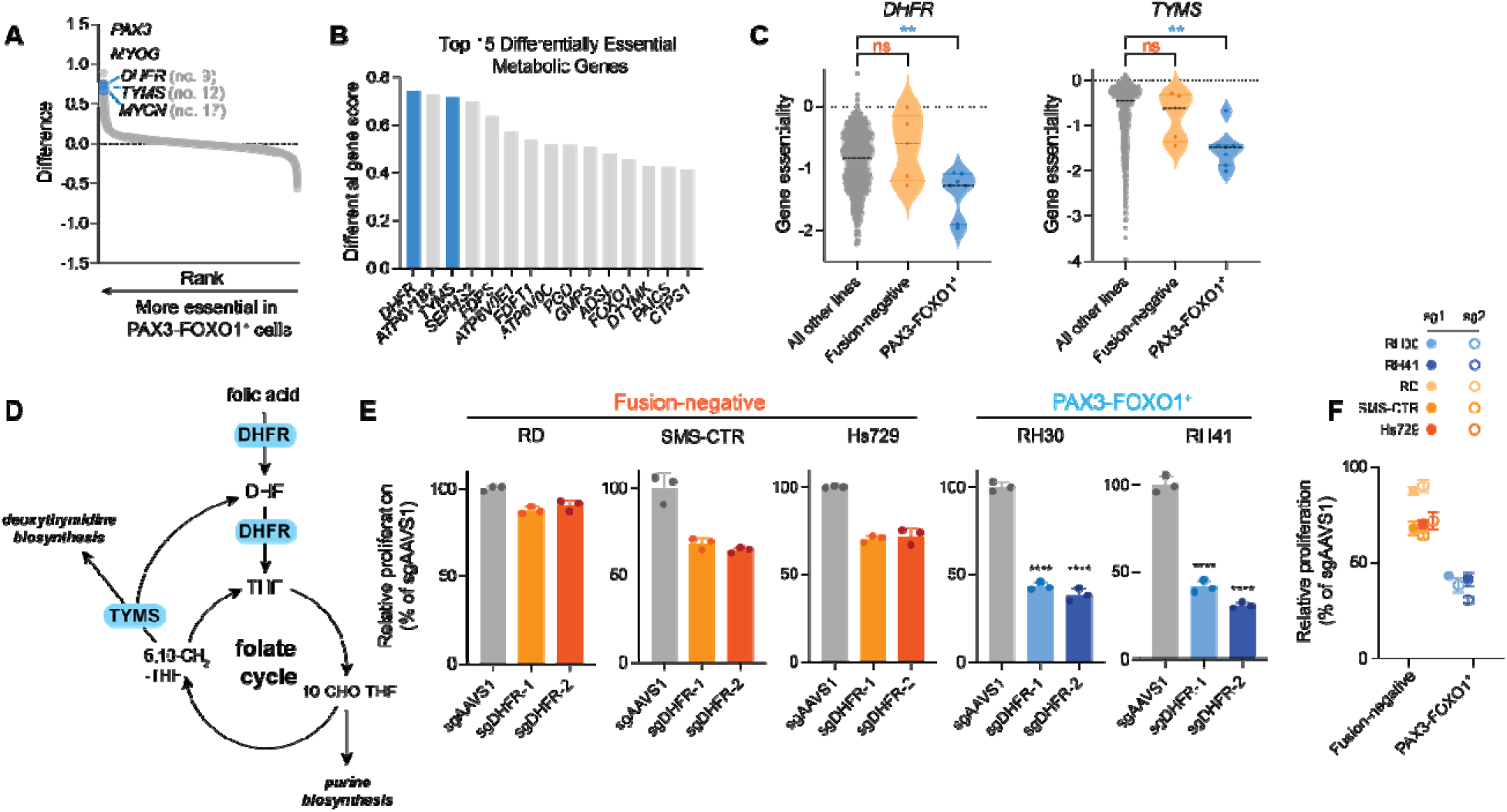
Genome-wide CRISPR screens reveal metabolic vulnerabilities specific to PAX3-FOXO1^+^ RMS. **A**, Results of differential essentiality analysis comparing PAX3-FOXO1^+^ and fusion negative cell lines. Genes are ranked in descending order of differential essentiality. Data were obtained from DepMap 24Q2 release^22^. **B**, Bar graph of the top 15 metabolic genes more essential in PAX3-FOXO1^+^ RMS cell lines compared to fusion-negative RMS cell lines. DHFR and TYMS are highlighted in blue. Data were obtained from DepMap 24Q2 release^22^. **C**, Violin plots of gene essentiality scores (obtained from DepMap 24Q2 release) for *DHFR* and *TYMS* in fusion-negative RMS cell lines (*n* = 5), PAX3-FOXO1^+^ RMS cell lines (*n* = 7), or all other cell lines (*n* = 1,121). Lines represent median of each group. **D**, Schematic depicting DHFR and TYMS in the folate cycle (DHF = dihydrofolate; THF = tetrahydrofolate; 5,10-CH_2_-THF = 5,10-methylene THF; 10-CHO-THF = 10-formyl THF). **E**, Relative proliferation of RMS cells in response to *DHFR* editing using two independent sgRNAs (sgDHFR-1, sgDHFR-2). Data are shown as a percentage of control (sgAAVS1) for each cell line, and are mean ± s.d., *n* = 3 independent replicates. **F**, Summary of effects of *DHFR* loss on proliferation in 3 fusion-negative and 2 PAX3-FOXO1^+^ RMS cell lines. Data are from the same experiment as data shown in **E**. Significance was assessed using one-way ANOVA with Dunnett’s (**C**) or Tukey’s (**E**) multiple comparisons test. In **C**, each RMS subtype was compared to all other lines in DepMap. In **E**, for each sgRNA, each PAX3-FOXO1^+^ cell line was compared to each fusion-negative cell line. Significance displayed is reflective of all comparisons. (\*\*\*\**P* < 0.0001)

To validate the differential dependence on DHFR/TYMS, we used CRISPR/Cas9 to engineer mutations in *DHFR* or *TYMS* in a panel of established RMS cell lines. To circumvent toxicity associated with loss of a common essential gene, cells were cultured with leucovorin calcium (a rescue agent for folate antagonists^32^) when edited with sgRNA targeting *DHFR* or thymidine and cytidine (rescue agents for impaired *de novo* pyrimidine synthesis) when edited with sgRNA targeting *TYMS* (Supplementary Fig. 2A). Following selection and confirmation of successful editing (Supplementary Fig. 2B, C), the rescue agent was withdrawn and proliferation was assessed. Across three fusion-negative cell lines, DHFR loss resulted in varying, yet modest, growth inhibition. In contrast, DHFR loss reduced proliferation by > 50% in two PAX3-FOXO1^+^ cell lines (Fig. 2E). Accordingly, across all sgRNA and cell lines used, DHFR deletion was more detrimental to PAX3-FOXO1^+^ cells than to their fusion-negative counterparts (Fig. 2F). Similar results were obtained following TYMS deletion in two RMS cell lines, although TYMS loss was overall less well-tolerated than DHFR loss (Supplementary Fig. 2D). Taken together, these data demonstrate that DHFR/TYMS are selective vulnerabilities of PAX3-FOXO1^+^ RMS cell lines.

### PAX3-FOXO1^+^ cells exhibit increased sensitivity to DHFR inhibition in vitro

Given the increased dependence of PAX3-FOXO1^+^ cells on DHFR, we asked whether inhibition of DHFR represents a specific and targetable vulnerability in these cells. Methotrexate (MTX) is a well-established inhibitor of DHFR and one of the oldest chemotherapeutic agents. Since its introduction in the 1960s, MTX has held an enduring role in the treatment of cancers including leukemia^33^ and osteosarcoma^34^, but its efficacy in PAX3-FOXO1^+^ RMS is largely unexplored. Previous, small-scale studies in RMS either occurred prior to recognition of the fusion oncogene or enrolled few fusion-positive patients^35–37^. Of note, the one confirmed PAX3-FOXO1^+^ patient examined exhibited stable disease after MTX treatment and then complete response to subsequent therapy^36^. We therefore aimed to determine the impact of MTX on RMS cells. Dose response curves revealed that PAX3-FOXO1^+^ RMS cells were more sensitive than fusion-negative RMS cells to doses of MTX ranging from 0.2 µM to 200 µM, the maximum dose tested (Fig. 3B). Consistently, whereas fusion-negative RMS cells maintained proliferation at approximately 50% of control rates despite the presence of 0.2 µM MTX, PAX3-FOXO1^+^ cells exhibited a more severe proliferative decline (RH30) or even an overall decrease in cell number (RH41) at this dose (Fig. 3C). Importantly, MTX sensitivity was not simply the result of different growth rates, as no consistent differences emerged in the basal proliferation rates of PAX3-FOXO1^+^ and fusion-negative cell lines (Supplementary Fig. 3A).

**Figure 3.**
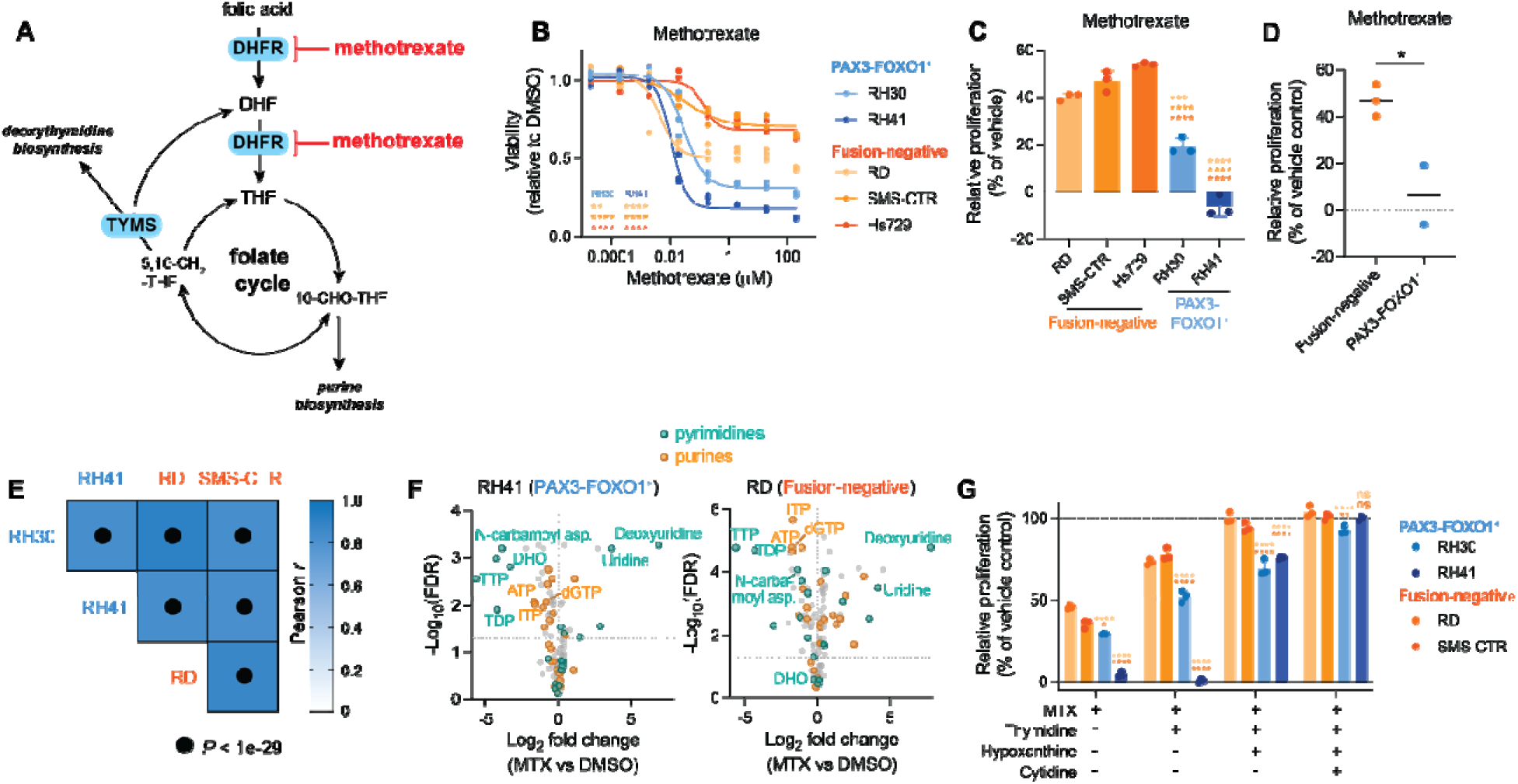
PAX3-FOXO1^+^ cells exhibit increased sensitivity to DHFR inhibition *in vitro*. **A**, Schematic of the folate cycle indicating targets of methotrexate (MTX). **B**, Dose response curve for MTX in RMS cell lines measured by CellTiter-Glo viability assay after 48 h of treatment. Maximum dose tested was 200 µM. **C**, Relative proliferation of RMS cells in response to MTX (0.2 µM) for 96 h. Data are shown as a percentage of the vehicle control (DMSO). **D**, Summary of the effects of MTX on relative proliferation from **C**. Each dot represents the average of all replicates from each cell line. **E**, Correlation matrix displaying Pearson correlation coefficients (*r*) for pairwise comparisons of metabolic effects of MTX in four RMS cell lines treated with MTX (0.2 µM) for 48 h. Correlation coefficients were calculated using the log_2_ fold change of MTX over vehicle (DMSO) for each metabolite. **F**, Volcano plots for two RMS cell lines (RH41, PAX3-FOXO1^+^ and RD, fusion-negative) showing the log_2_ fold change in metabolite abundance in cells treated with MTX (0.2 µM, 48 h) compared to cells treated with vehicle (DMSO). Dotted horizontal line indicates significance threshold FDR < 0.05. **G**, Relative proliferation of RMS cells in response to MTX (0.2 µM) in the presence or absence of thymidine (10 µM), hypoxanthine (100 µM), and/or cytidine (10 µM) for 96 h. Data are shown as a percentage of the vehicle control (DMSO, H_2_O, 0.1N NaOH). For **B**, **C**, and **G**, data are mean ± s.d., *n* = 3 independent replicates. Significance was assessed by ordinary one-way ANOVA with Tukey’s multiple comparisons test (**B**, **C**) and unpaired two-tailed Student’s *t*-test (**G**). In **B**, **C**, and **G**, each PAX3-FOXO1^+^ cell line was compared to each fusion-negative cell line. For **B**, comparison shown at the 0.2 µM dose. (ns = not significant, \**P* < 0.05, \*\**P* < 0.01, \*\*\**P* < 0.001, \*\*\*\**P* < 0.0001)

DHFR executes two reactions: the sequential reduction of folic acid to DHF and then to THF (Fig. 3A). Although folic acid is present in most standard cell culture media, 5-methyl tetrahydrofolate (5-meTHF) is the primary physiological source of folate *in vivo*^38,39^. To test the possibility that MTX sensitivity was an artifact of the presence of folic acid, we replaced folic acid with 5-meTHF. As expected based on previous results^38,39^, cells with 5-meTHF as their primary folate source were overall less sensitive to MTX (Supplementary Fig. 3B). Nevertheless, PAX3-FOXO1^+^ cells still exhibit greater sensitivity to MTX than fusion-negative RMS cells, even with 5-meTHF as the primary folate source. (Supplementary Fig. 3B). These results suggest that PAX3-FOXO1^+^ cells require DHFR for purposes beyond provision of reduced folates.

To further test the notion that PAX3-FOXO1^+^ cells are preferentially reliant on DHFR, but not folate metabolism more generally, we tested the impact of other drugs targeting the folate cycle. Neither pemetrexed, which inhibits multiple steps of the folate cycle, nor SHIN2, which targets serine hydroxymethyltransferase (SHMT) 1/2, exhibited differential efficacy in fusion-negative and PAX3-FOXO1^+^ RMS cells (Supplementary Fig. 3D, G, J). Of the many fates of the one-carbon units carried by the folate cycle, *de novo* purine biosynthesis is thought to represent the largest demand^40^; accordingly, MTX is frequently considered to act as an anti-purine^41,42^. To assess the impact of purine synthesis inhibition, we treated cells with lometrexol, a phosphoribosylglycinamide formyltransferase (GART) inhibitor. Notably, lometrexol had equivalent effects on viability and proliferation in PAX3-FOXO1^+^ and fusion-negative RMS cells, indicating that increased demand for *de novo* purine synthesis does not drive enhanced sensitivity to MTX in PAX3-FOXO1^+^ RMS cells (Supplementary Fig. 3E, H, K). More generally, actinomycin D and cyclophosphamide are currently deployed in standard-of-care chemotherapeutic regimens for RMS patients^13^, but PAX3-FOXO1^+^ and fusion-negative cells did not exhibit differential sensitivity to either actinomycin D or 4-hydroperoxy cyclophosphamide (4-HC), the active metabolite of cyclophosphamide (Supplementary Fig. 3F, I, L). Collectively, these data support the notion that, of all drugs evaluated, differential drug sensitivity between RMS subtypes is unique to DHFR inhibition via MTX.

### Demand for nucleotides underlies methotrexate sensitivity

We next asked why PAX3-FOXO1^+^ cells are more sensitive to MTX than their fusion-negative counterparts. To this end, we assessed the metabolic consequences of MTX treatment in PAX3-FOXO1^+^ and fusion-negative RMS cell lines. To our surprise, the metabolic response to MTX was largely independent of RMS subtype: across all 4 cell lines, the metabolic changes induced by MTX were highly correlated (Pearson *r* > 0.9) (Fig. 3E). These results demonstrate that MTX exerts equivalent effects in all cell lines, and thus differential MTX sensitivity is not due to changes in drug uptake or metabolism.

We therefore examined individual metabolites to determine which limit proliferation in MTX-treated cells. Consistent with the noted anti-purine effects of MTX^41,42^, purine nucleotides including adenosine triphosphate (ATP), inosine triphosphate (ITP), and deoxyguanosine triphosphate (dGTP) were consistently decreased across cell lines (Fig. 3F, Supplementary Fig. 3M). However, hypoxanthine, a purine salvage precursor that mitigates MTX toxicity in certain circumstances^43,44^, failed to rescue growth in the presence of MTX, and even made one cell line (RH30) noticeably more sensitive to MTX (Supplementary Fig. 3N). These results are consistent with the observations that folate cycle and purine biosynthesis inhibitors are equivalently effective across RMS cell lines and argue that MTX sensitivity in PAX3-FOXO1^+^ cells is not the result of increased demand for *de novo* purine biosynthesis.

We therefore tested the hypothesis that differential sensitivity to MTX arises from its inhibition of *de novo* thymidine synthesis. MTX been shown to interfere with *de novo* thymidine synthesis via inhibition of TYMS^45,46^, and thymidine supplementation can rescue certain cells from MTX^47,48^. Indeed, MTX strongly depleted pyrimidine nucleotides thymidine diphosphate (TDP) and thymidine triphosphate (TTP) as well as pyrimidine synthesis intermediates N-carbamoyl aspartate and dihydroorotate (Fig. 3F, Supplementary Fig. 3M). Thymidine supplementation alone proved effective at rescuing fusion-negative cell lines from MTX, restoring proliferation to 73% (RD) and 77% (SMS-CTR) of untreated cells (Fig. 3G). In contrast, thymidine had a more modest (RH30, 52%) or no (RH41) ability to rescue PAX3-FOXO1^+^ cells from MTX (Fig. 3G). Whereas hypoxanthine had little effect on its own, co-treatment with hypoxanthine and thymidine fully rescued proliferation in fusion-negative cells, demonstrating that thymidine synthesis is the most acute limitation for these cells, and combined restoration of thymidine and purine synthesis restores proliferation of fusion-negative RMS cells in the presence of MTX (Fig. 3G). In contrast, PAX3-FOXO1^+^ cells maintained a significant proliferative defect despite improved proliferation in the presence of hypoxanthine and thymidine (Fig. 3G). Given that MTX also reduced levels of intermediates involved in *de novo* pyrimidine synthesis (Fig. 3F, Supplementary Fig. 3M), we further supplemented cells with the remaining pyrimidine nucleoside, cytidine. The combined addition of thymidine, cytidine, and hypoxanthine fully restored proliferation of MTX-treated PAX3-FOXO1^+^ cells back to control levels (Fig. 3G). Collectively, these observations indicate that MTX impairs proliferation by limiting nucleotide synthesis, as restoring nucleotides is sufficient to reverse growth inhibition in the presence of MTX. Furthermore, the requirement for both cytidine and thymidine to rescue PAX3-FOXO1^+^ cells, but not fusion-negative RMS cells, reveals that increased demand for pyrimidine nucleotides contributes to MTX sensitivity in PAX3-FOXO1^+^ cell lines.

### Methotrexate phenocopies loss of PAX3-FOXO1

The observation that demand for pyrimidine nucleotides drives differential sensitivity to MTX in RMS cell lines led us to hypothesize that MTX could be used to target PAX3-FOXO1-driven cell states. Suggestively, *PAX3-FOXO1* silencing, which is associated with downregulation of genes associated with the cycling progenitor cell state, also drives downregulation of pyrimidine-related genes and reduced expression of both DHFR and TYMS (Supplementary Fig. 4A, B). We therefore compared the metabolic response of *PAX3-FOXO1* silencing to that of MTX treatment in RH30 cells. Principal component analysis revealed that cells treated with MTX or in which *PAX3-FOXO1* was silenced clustered together, away from their respective controls, suggesting underlying similarities in the metabolic effects of MTX and *PAX3-FOXO1* silencing (Fig. 4A). Indeed, across all profiled metabolites, the changes induced by *PAX3-FOXO1* silencing significantly correlated (*r* = 0.5351, *P* < 3.29e-8) with changes induced by MTX (Fig. 4B). The correlation persisted even when only pyrimidine-related metabolites were assessed, driven in part by the coordinated downregulation of N-carbamoyl aspartate, dihydroorotate, TDP, and TTP following both MTX treatment and *PAX3-FOXO1* silencing (Fig. 4B). Thus, MTX treatment and *PAX3-FOXO1* silencing exert similar effects on cell metabolism—and, in particular, on pyrimidine metabolism.

**Figure 4.**
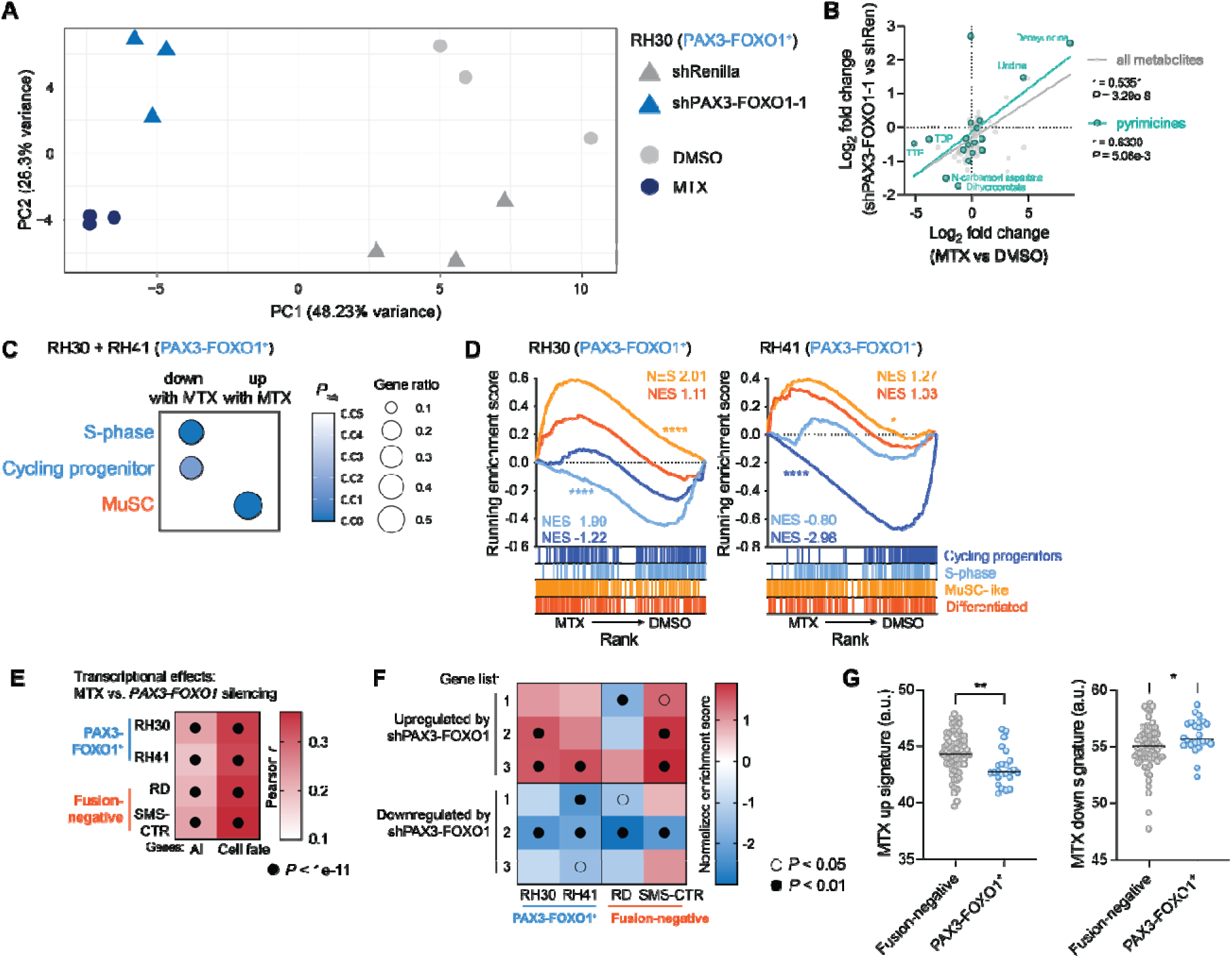
Methotrexate phenocopies loss of PAX3-FOXO1. **A**, Principal component analysis of RH30 cells treated with vehicle (DMSO) or MTX (0.2 µM) for 48 h, and RH30 cells expressing shRNA against *Renilla* luciferase (control) or *PAX3-FOXO1* (cultured with doxycycline for 72 h to induce knockdown). MTX data were previously shown in Fig. 1A and shPAX3-FOXO1 data were previously shown in Fig. 1D. 101 metabolites were included in analysis. **B**, Scatter plot of metabolic data shown in **A** comparing metabolic effects of MTX and *PAX3-FOXO1* silencing in RH30 cells. Pyrimidine metabolites are highlighted in teal. **C**, Overrepresentation analysis of genes significantly (*P_adj_* < 0.05) upregulated (“up with MTX”) or downregulated (“down with MTX”) upon MTX treatment in two PAX3-FOXO1^+^ cell lines (RH30, RH41). Analysis was performed using cell fate gene lists detailed in Supplementary Table 3. Cells were treated with MTX (0.2 µM) for 48 h. **D**, Gene set enrichment analysis (GSEA) depicting the effect of MTX on cell fate gene lists in two PAX3-FOXO1^+^ cell lines (RH30, RH41). Cells were treated with MTX (0.2 µM) for 48 h. **E**, Heatmap of Pearson correlation coefficients (*r*) comparing transcriptional effects of MTX (in each RMS cell line) against effects of *PAX3-FOXO1* silencing (in KFR cell line, published data^20^). Correlation coefficients were calculated for all genes and for genes involved in cell fate. Cells were treated with MTX (0.2 µM) for 48 h. **F**, Heatmap depicting normalized enrichment scores from GSEA analysis of data from four MTX-treated RMS cell lines. Gene lists (1, 2, and 3) are composed of genes significantly upregulated or downregulated by PAX3-FOXO1 in three independent published experiments^20,49^. **G**, Gene set signature scores in published RNA-sequencing of RMS patient tumors^50^ using gene lists composed of genes significantly (*P* < 0.05) upregulated (“MTX up”) or downregulated (“MTX down”) by MTX in two PAX3-FOXO1^+^ cell lines (RH30, RH41). Gene lists are provided in Supplementary Table 3. Significance was assessed in **G** by unpaired two-tailed Student’s *t* test comparing RMS subtypes. (\**P* < 0.05, \*\**P* < 0.01, \*\*\*\**P* < 0.0001)

To determine whether the metabolic changes accompany changes in cell state, we performed RNA-sequencing of cells exposed to MTX for 48 h. Genes defining the PAX3-FOXO1-driven cycling progenitor and S-phase cell states were significantly (*P* < 0.05) overrepresented among genes decreased by MTX in both RH30 and RH41 cells (Fig. 4C). Reciprocally, genes defining the fusion-negative associated MuSC cell state were significantly overrepresented among genes upregulated by MTX in both PAX3-FOXO1^+^ cell lines (Fig. 4C). More broadly, gene set enrichment analysis of each cell line’s response to MTX revealed that genes downregulated by MTX were significantly enriched (*P* < 0.05) for genes defining the cycling progenitor cell state whereas genes defining the fusion-negative MuSC cell state were enriched among genes upregulated by MTX in PAX3-FOXO1^+^ cells (Fig. 4D). Similar trends were observed in fusion-negative cells, with cycling progenitor genes enriched among genes downregulated by MTX, although the effect of MTX on gene expression programs was less consistent than in PAX3-FOXO1^+^ cells (Supplementary Fig. 4C). Importantly, the proportion of cells in S-phase was largely consistent between PAX3-FOXO1^+^ and fusion-negative cell lines, either in untreated or MTX-treated cells, as measured by 5-ethynyl-2′-deoxyuridine (EdU) incorporation (Supplementary Fig. 4D). Together, these results indicate that MTX alters gene programs associated with RMS cell state, particularly in PAX3-FOXO1^+^ cells, without exerting differential effects on cell cycle distribution in the different RMS subtypes.

The observation that MTX and *PAX3-FOXO1* silencing induce similar metabolic alterations led us to ask whether MTX phenocopies the impact of *PAX3-FOXO1* silencing on transcriptional networks. To this end, we quantified the impact of *PAX3-FOXO1* silencing on all genes expressed in a PAX3-FOXO1^+^ cell line and compared these changes with those induced by MTX in four RMS cell lines. Across all genes in the genome, changes induced by MTX significantly correlated with changes induced by *PAX3-FOXO1* silencing. When focusing on genes related to RMS cell states, the changes induced by MTX and *PAX3-FOXO1* silencing were even more correlated, demonstrating that MTX and *PAX3-FOXO1* silencing exert similar impact on genes related to RMS cell state (Fig. 4E). More generally, genes identified as up- or down-regulated across three separate experiments assessing the impact of *PAX3-FOXO1* silencing^20,49^ were significantly enriched among genes up- and down-regulated, respectively, following MTX treatment (Fig. 4F). To examine whether MTX-regulated genes are altered in human tumors, we analyzed expression of genes reproducibly up- or down-regulated by MTX across PAX3-FOXO1^+^ RMS cell lines in culture in datasets generated from fusion-negative and PAX3-FOXO1^+^ human RMS tumors^50^. These results revealed that genes repressed following MTX treatment tended to be expressed at higher levels in PAX3-FOXO1^+^ tumors whereas genes induced by MTX tended to be expressed at lower levels in PAX3-FOXO1^+^ tumors (Fig. 4G). Taken together, these results demonstrate that MTX mimics the transcriptional and metabolic effects of PAX3-FOXO1 inhibition, supporting the notion that MTX targets PAX3-FOXO1-dependent cell states.

### Methotrexate is effective as a single agent in PAX3-FOXO1^+^ tumors

The observation that MTX phenocopies *PAX3-FOXO1* silencing raises the possibility that MTX could be used to target PAX3-FOXO1-driven tumors. To investigate how RMS tumors respond to MTX *in vivo*, we xenografted established RMS cell lines into the flanks of nude mice. Once tumors were established and reached 100 mm^3^ in size, mice were treated with 30 mg/kg MTX three times a week, a dose approximately 10-fold lower than those mimicking pediatric high-dose MTX regimens in mice (400 mg/kg)^51^. 24 h after each dose, mice were provided leucovorin calcium, a rescue agent employed clinically to mitigate toxicity following MTX treatment (Fig. 5A). This dose regimen of MTX was remarkably well-tolerated, and mouse weights remained relatively constant throughout the duration of treatment in both vehicle- and MTX-treated groups (Supplementary Fig. 5A, B). Neither fusion-negative xenograft (RD and SMS-CTR) displayed a robust response to MTX, and MTX neither slowed tumor growth rate (Fig. 5B, F) nor significantly affected survival of mice engrafted with these cell lines (Fig. 5D). In contrast, both xenografts derived from PAX3-FOXO1^+^ cell lines (RH30 and RH41) showed a striking response to MTX, with tumor growth noticeably slower in MTX-treated mice (Fig. 5C, G). Accordingly, MTX was sufficient to significantly prolong survival of mice bearing PAX3-FOXO1^+^ xenografts when compared to matched vehicle controls, even when provided as a single agent (Fig. 5E). While tumors treated with MTX eventually escaped the growth-inhibitory effects of MTX, these results suggest that MTX could be an effective addition to existing chemotherapeutic regimens specifically for the treatment of PAX3-FOXO1^+^ RMS.

**Figure 5.**
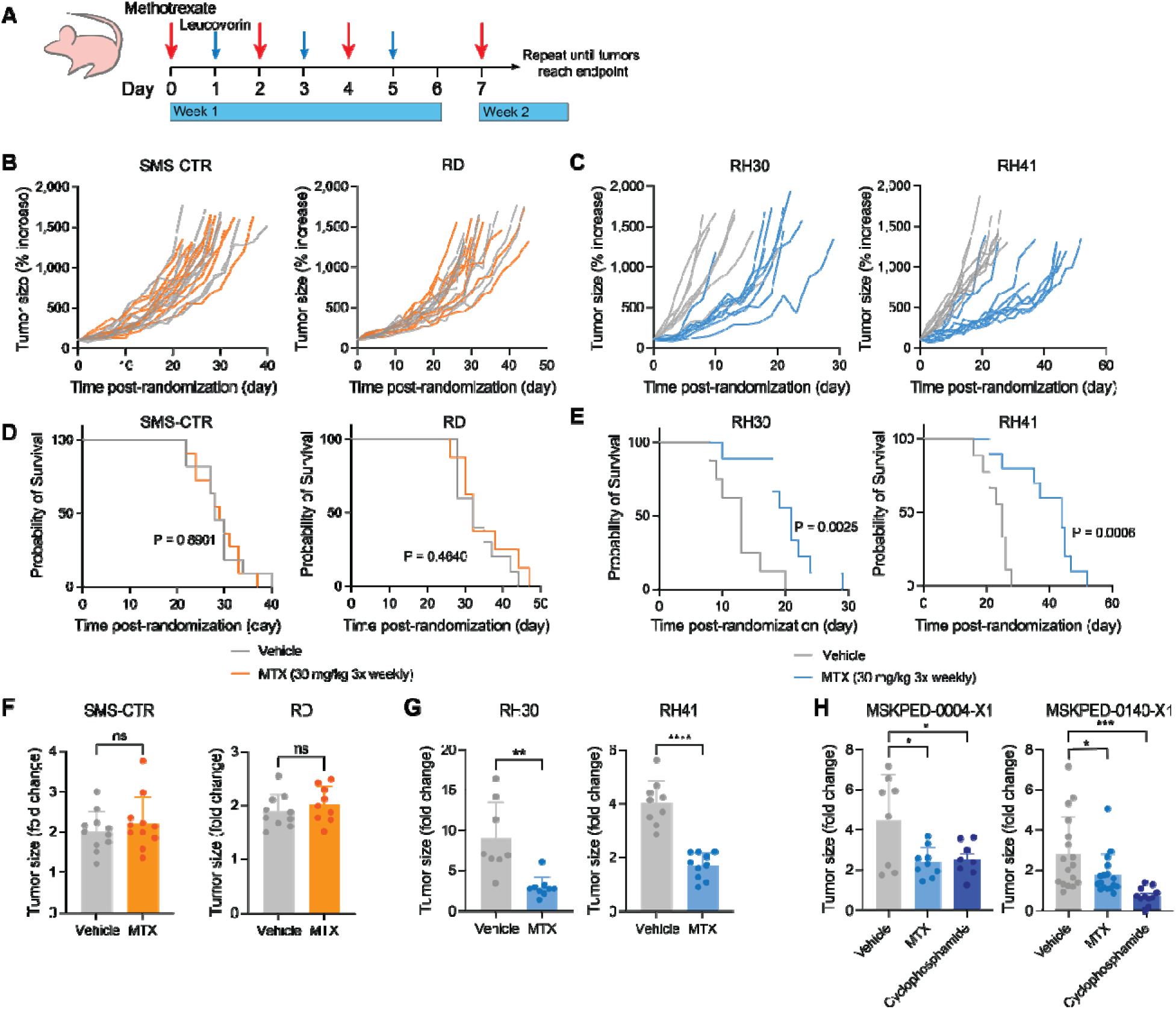
Methotrexate is effective as a single agent in PAX3-FOXO1^+^ tumors. **A**, Schematic depicting dosing strategy for methotrexate and leucovorin rescue in RMS cell line xenograft experiments. **B**, **C**, Tumor growth curves for fusion-negative RMS (**B**, SMS-CTR and RD) and PAX3-FOXO1^+^ RMS (**C**, RH30 and RH41) cell line xenografts. Data are normalized to tumor size at the start of treatment. **D**, **E**, Survival curves for mice bearing fusion-negative RMS (**D**, SMS-CTR and RD) or PAX3-FOXO1^+^ RMS (**E**, RH30 and RH41) cell line xenografts. **F**, **G**, Change in tumor size for fusion-negative RMS (**F**, SMS-CTR and RD) and PAX3-FOXO1^+^ RMS (**G**, RH30 and RH41) cell line xenografts over the first week (6-8 d) of treatment with vehicle (corn oil) or MTX. Data are shown as a fold change relative to tumor size at the start of treatment. **H**, Change in tumor size for two PAX3-FOXO1^+^ PDX models (MSKPED-0004-X1, left and MSKPED-0140-X1, right) over the first week of treatment with vehicle (corn oil), MTX, or cyclophosphamide. Data are shown as a fold change relative to tumor size at the start of treatment. Data are mean ± s.d. For SMS-CTR (**B**, **D**, **F**), *n* = 11. For RD (**B**, **D**, **F**), *n* = 10 (vehicle) or 9 (MTX). For RH30 (**C**, **E**, **G**), *n* = 8 (vehicle) or 9 (MTX). For RH41 (**C**, **E**, **G**), *n* = 9 (vehicle) or 10 (MTX). For MSKPED-0004-X1 (**H**), *n* = 8 (vehicle, cyclophosphamide) or 9 (MTX). For MSKPED-0140-X1 (**H**), *n* = 17 (vehicle, MTX) or 11 (cyclophosphamide). Significance was assessed by log-rank (Mantel-Cox) test (**C**, **E**), unpaired two-tailed Student’s *t*-test (**F**, **G**), or ordinary one-way ANOVA with Dunnett’s multiple comparisons test (**H**). In **H**, each treatment group was compared to vehicle control. (ns = not significant, \**P* < 0.05, \*\**P* < 0.01, \*\*\**P* < 0.001, \*\*\*\**P* < 0.0001)

We next evaluated the efficacy of MTX in two patient-derived xenograft (PDX) models of PAX3-FOXO1^+^ RMS. Of note, cyclophosphamide, currently employed in standard-of-care regimens, was effective at slowing tumor growth and prolonging survival in only one of two examined PAX3-FOXO1^+^ PDX models (Supplementary Fig. 5C, D). Nevertheless, MTX was effective in significantly slowing tumor growth during the first week of treatment in both PAX3-FOXO1^+^ PDX models (Fig. 5H), including a highly aggressive and chemoresistant model (MSKPED-0004-X1). When leucovorin administration was appropriately delayed, MTX was sufficient to prolong survival as a single chemotherapeutic agent (Supplementary Fig. 5D). In the model where leucovorin was provided continuously after an initial 24 h delay (Supplementary Fig. 5C), MTX was initially effective at slowing growth, but this effect was not durable. While there is ample scope to improve dosing strategies, these data indicate that MTX treatment is able to slow tumor growth—and even increase survival—in models of PAX3-FOXO1^+^ RMS.

## Discussion

Here we show that PAX3-FOXO1^+^ RMS cells exhibit metabolic dependence on pyrimidine synthesis that can be exploited with MTX. New therapies are urgently needed for patients with PAX3-FOXO1^+^ tumors, as these patients have an especially grim prognosis. Moreover, current aggressive treatment regimens, even when successful, are associated with significant risk for lifelong morbidities in pediatric patients^52^. As targeting transcription factors is notoriously difficult, our work suggests that targeting metabolic vulnerabilities inherent to the PAX3-FOXO1-driven cell state represents an alternative and tractable strategy for treating tumors driven by this fusion oncogene. MTX has the additional advantage of being well-tolerated clinically: leucovorin enables safe administration of high doses, even in pediatric patients^32,53^, such that over a course of 10 years, less than two percent of pediatric patients receiving high-dose MTX required intense supportive care^54^. Furthermore, the dose of 30 mg/kg utilized in this study for *in vivo* experiments is approximately equivalent to a dose of 90 mg/m^2^ in humans^55^, appreciably lower than the 500 mg/m^2^ that is considered the minimum for high-dose MTX^32^. Indeed, MTX doses for osteosarcoma patients often range from 8-12 g/m^2^ ^34^, over 100-fold higher than the doses used in this study. The potential to leverage a relatively low dose of MTX for PAX3-FOXO1^+^ RMS is appealing given that this disease predominantly strikes pediatric patients and provides ample scope for combination with existing chemotherapies, possibly allowing for the administration of lower and less toxic doses of the current standard-of-care drugs. Further optimization of the delivery and dosing schedule for MTX—with accompanying leucovorin rescue—and potential for combination with existing therapeutic options will be a critical direction for future investigation.

Our data indicate that MTX specifically targets the PAX3-FOXO1-driven malignant cell state. Although PAX3-FOXO1 does not directly activate genes involved in pyrimidine biosynthesis, expression of these genes depends on sustained PAX3-FOXO1 expression and PAX3-FOXO1 loss reduces metabolites associated with pyrimidine biosynthesis. Our results are consistent with the notion that PAX3-FOXO1 induces a metabolic addiction, akin to oncogene addiction, in which the specific metabolic program directed by PAX3-FOXO1 (pyrimidine synthesis) is required for survival of PAX3-FOXO1^+^ cells. Accordingly, targeting pyrimidine synthesis with MTX mimics loss of PAX3-FOXO1, recapitulating the metabolic and transcriptional effects of PAX3-FOXO1 silencing. Intriguingly, the induction of a DHFR-dependent metabolic state may be a general principle of PAX3 activation. During development, PAX3 plays a critical role in the formation of skeletal muscle and the central nervous system^56^, and *Pax3* mutant mice exhibit neural tube defects that can be mitigated by supplementation of folic acid but not formate^57^. These observations indicate that defects in DHFR activity specifically, but not one-carbon metabolism broadly, underlie developmental disorders driven by reduced PAX3 function and suggest that PAX3-FOXO1^+^ malignant cell states mimic aspects of PAX3-directed cell states during normal development. Determining why PAX3-driven cell states have increased demand for DHFR activity—whether because of increased demand for pyrimidine nucleotides, reduced ability to salvage pyrimidine nucleotides, or general sensitivity to nucleotide imbalance—will provide important insight into additional strategies to treat diseases driven by aberrant PAX3 transcriptional activity.

More broadly, this work reinforces the notion that identifying metabolic dependencies of distinct cell states will provide opportunities to refine therapeutic approaches for pediatric solid tumors like RMS. Although the metabolic landscape of pediatric solid tumors like RMS is still relatively unexplored, the concept of targeting metabolism for cancer treatment is as old as chemotherapy itself. MTX, one of the first antimetabolites, has been continuously used for over sixty years to treat various cancers, both as a frontline and relapse therapy^58^. However, like all traditional chemotherapies, it has not had universal success, highlighting the need to identify specific disease subtypes and cell states in which MTX will be most effective. Continued exploration of metabolic dependencies of malignant cell states is likely to provide new avenues to treat pediatric cancers, where low mutational burden, especially in fusion-positive RMS where PAX3-FOXO1 (or alternate fusion PAX7-FOXO1) is itself the main oncogenic driver, presents a significant challenge to targeted therapy development^50^. Alternative approaches exploiting the metabolic rewiring necessary for maintenance of aberrant cell fate programs may result in unique metabolic vulnerabilities that can be exploited for therapeutic benefit.

## Materials & Methods

### Cell Culture

RH30 (CRL-2061), RD (CCL-136), and Hs729 (HTB-153) were purchased from the ATCC. RH41 and SMS-CTR were obtained from the Children’s Oncology Group Childhood Cancer Repository. Cell lines were maintained in DMEM supplemented with 10% fetal bovine serum (Gemini). All cells were routinely tested for mycoplasma.

### Xenograft studies

All animal work was approved by Memorial Sloan Kettering Cancer Center (MSKCC) Institutional Animal Care and Use Committee (protocol nos. 19-09-014, Finley and 16-08-011, Kung) in compliance with all relevant ethical regulations. Animals were housed in a pathogen-free facility maintained at 21.5L±L1L°C and relative humidity 30–70% under a 12/12 h light/dark cycle.

For cell line xenograft experiments, 1 x 10^6^ (RH30), 2 x 10^6^ (RD and SMS-CTR), or 4 x 10^6^ (RH41) cells were mixed 1:1 with Matrigel (Corning, 356231) and subcutaneously engrafted into the right flank of 6– 8-week-old female athymic nude mice (Charles River Laboratories, strain code 490). Mice were enrolled in experimental arms when their tumor size reached 100 mm^3^. Due to the staggered enrollment of mice in each study, female mice were exclusively used to avoid single housing of mice. Following enrollment, mice were treated with methotrexate (Sigma, M9929, 30 mg/kg) dissolved in corn oil (Sigma, C8267) and administered via IP injection three times a week, with each dose followed 24 h later by an IP injection of leucovorin calcium (Sigma, PHR1541, 24 mg/kg) dissolved in saline solution. Vehicle controlled mice received an equivalent volume of corn oil three times a week. Tumors were measured with calipers three times a week, and tumor volume was calculated using the equation: volume = 0.5 × length × width^2^. Mice were euthanized once tumors reached endpoint criteria (size at or exceeding 1500 mm^3^), or if mandated by veterinary staff. Mouse weight was recorded at time of enrollment and at least once a week once mice were enrolled into each study.

For patient-derived xenograft experiments, NSG mice (Jackson Labs, strain #005557) were anesthetized and sterilized with 70% ethanol. Viably cryopreserved tumor fragments measuring 1-2 mm were thawed in a 37°C water bath and then implanted into a subcutaneous pocket in the mouse flank generated by blunt dissection. Surgical incisions were approximated and sealed with Vetbond tissue adhesive (MWI Veterinary Supply, 006245). An equal number of male and female mice were implanted. Implanted mice were monitored closely for tumor engraftment and growth. Mice were enrolled in treatment arms once tumors reached approximately 100 mm^3^ in size, balancing the numbers of male and female mice in each group as much as possible. For the experiment utilizing the model MSKPED-0140-X1, mice were treated with methotrexate (with leucovorin rescue) following a regimen identical to that used for cell line xenograft experiments (see above). For the experiment utilizing the model MSKPED-0004-X1, mice were treated with methotrexate (30 mg/kg) administered three times a week. 24 h after the first dose of MTX was administered, leucovorin calcium was provided continuously in drinking water at a concentration of 100 mg/L. Medicated water was refreshed twice a week. Cyclophosphamide (Sigma, C0768, 50 mg/kg) was administered to mice via IP injection once daily for five consecutive days.

### Generation of CRISPR/Cas9-edited lines

Single guide RNA (sgRNA) sequences targeting *DHFR*, *TYMS* or the AAVS1 safe harbor site (control) were cloned into lentiCRISPRv2 (Addgene, 52961) as previously described^59^ but using BsmbI (Thermo Fisher Scientific, FD0454) as the restriction enzyme. All sgRNA sequences are provided in Supplementary Table 2. Lentivirus was produced and introduced into cells as described below, followed by selection with 1 μg/mL (RH30, RD, SMS-CTR, Hs729) or 0.5 μg/mL (RH41) puromycin (Thermo Fisher Scientific, A1113803). sgDHFR cell lines were maintained in the continuous presence of 100 μM leucovorin calcium (Fisher Scientific, 18-603-788) until plated for experiments. sgTYMS cell lines were maintained in the continuous presence of 100 μM thymidine (Sigma, T9250) and 100 μM cytidine (Sigma, C4654) until plated for experiments.

### Generation of shPAX3-FOXO1 lines

Due to the existence of an EcoRI site at the PAX3-FOXO1 junction, the canonical miR-E hairpin cloning protocol^60^ was modified as follows. Hairpin sequences targeting the fusion junction between PAX3 and FOXO1 were designed using splashRNA^61^ and additional bases were added to the beginning and end of the sequence to enable PCR amplification. All shRNA sequences and cloning primers are provided in Supplementary Table 2. Hairpins were PCR-amplified and the doxycycline-inducible lentiviral vector LT3GEPIR (ref^60^) (Addgene, 111177) was digested with EcoRI and XhoI. A hairpin sequence targeting *Renilla* luciferase was used as a control and was cloned into LT3GEPIR following the standard miR-E hairpin cloning protocol^60^. For each hairpin, the PCR-amplified insert and digested vector were assembled using NEBuilder HiFi DNA Assembly Master Mix (New England Biolabs, E2621S). A hairpin sequence targeting *Renilla* luciferase was used as a control and was cloned into LT3GEPIR following the standard miR-E hairpin cloning protocol^60^. Lentivirus was produced and introduced into RH30 cells as described below, followed by selection with 1 μg/mL puromycin (Thermo Fisher Scientific, A1113803).

### Lentiviral Production

Lentivirus was generated by co-transfection of lentiviral vectors expressing sgRNA or shRNA of interest with packaging plasmids psPAX2 (Addgene, 12260) and pMD2.G (Addgene, 12259) into 293T cells via calcium phosphate transfection. Viral-containing supernatant was passed through a 0.45 μm filter to remove debris. Target cells were exposed to viral supernatants in the presence of polybrene (8 μg/mL, Sigma, 107689) for two 24 h periods in 6-well plates before being moved to 10 cm^2^ dishes. Antibiotic selection was started 24 h later.

### Growth curves

To measure response to drug treatment, cells were seeded into 12-well plates and allowed to adhere overnight. The following day, three wells of each line were counted to measure the starting cell number. If applicable, the remaining cells were changed to the relevant treatment medium. Cells were counted 96 h later using a Beckman Coulter Multisizer 4e with a cell volume gate of 400-20,000 fL. Cell counts were normalized to starting cell number, and the number of population doublings per day was calculated using the following equation:

Doublings per day = [log_2_(final cell count/initial cell count)]/4

Relative proliferation for each treatment group is expressed as a percentage of the control group (set to 100%). This percentage was calculated using the doublings per day for each group. Methotrexate (Sigma, M9929), pemetrexed (Sigma, SML1490), lometrexol (Cayman Chemical Company, 18049), actinomycin D (Sigma, A1410), and 4-Hydroperoxy cyclophosphamide (MedChemExpress, HY-117433) were used at indicated concentrations. SHIN2 (ref^62^) was a generous gift from the Rabinowitz lab. For nucleobase or nucleoside rescue experiments, thymidine (Sigma, T9250), cytidine (Sigma, C4654), and/or hypoxanthine (Sigma, H9377) were added at stated concentrations. To measure response to leucovorin withdrawal in sgDHFR lines, cell lines were seeded into 12-well plates in media without leucovorin calcium. The following day, three wells of each line were counted to measure the starting cell number. Media was refreshed two days later, and cells were counted five days after the starting cell number was measured. Cell counts were normalized to starting cell number, and the equation above was used to calculate doublings per day but with 5 as the denominator. The percentage of cells in each knockout line relative to the control line (sgAAVS1) was then calculated. To measure response to thymidine and cytidine withdrawal in sgTYMS lines, a similar procedure to that described above for leucovorin withdrawal was used, with two key modifications: cell lines were seed into 12-well plates without thymidine or cytidine, and cells were counted 4 days after the starting cell number was measured. For evaluation of the effects of PAX3-FOXO1 knockdown, cells were seeded as described above and doxycycline (1 μg/mL, Sigma, D9891) was added 24 h later. Cells were counted every day for three days. Medium containing doxycycline was refreshed every 48 h.

### Viability Assay

Cells were seeded into clear-bottom, white 96-well plates (Corning, 3610) and allowed to adhere overnight. Serial drug dilutions were added 24 h later, and the CellTiter-Glo viability assay (Promega, G7572) was run 48 h later following manufacturer instructions. For evaluating response to MTX with 5-meTHF as the folate source, cells were seeded into 96-well plates in DMEM without folic acid (MSKCC media core) supplemented with 10% fetal bovine serum and 4 mg/L 5-meTHF (Sigma, M0132).

### LC-MS Analysis

Cells were seeded in six-well plates. If applicable, 24 h after seeding, cells were changed to treatment media containing vehicle control (DMSO, Sigma, D2650) or 0.2 μM methotrexate (Sigma, M9929), or 1 μg/mL doxycycline (Sigma, D9891). 48 h later, cells were harvested using 1 mL ice-cold 80% methanol containing 2 μM deuterated 2-hydroxyglutarate (d-2-hydroxyglutaric-2,3,3,4,4-d5 acid (d5-2HG)) as an internal standard. Plates were incubated at -80°C overnight. Samples were centrifuged at 20,000 x *g* for 20 min at 4°C to clear protein, then dried in an evaporator (Genevac EZ-2 Elite). Dried extracts were resuspended in 30 μL of mobile phase A (10 mM tributylamine and 15 mM acetic acid in 97:3 water:methanol) and were incubated on ice for 20 min with vortexing every 5 min. Samples were cleared by centrifugation at 20,000 x *g* for 20 min at 4°C. After clearing, samples were subjected to MS/MS acquisition using an Agilent 1290 Infinity LC system equipped with a quaternary pump, multisampler, and thermostatted column compartment coupled to an Agilent 6470 series triple quadrupole system with a dual Agilent Jet Stream source for sample introduction. Data were acquired in dynamic MRM mode using electrospray ionization in negative ion mode. Capillary voltage was 2000 V, nebulizer gas pressure was 45 Psi, drying gas temperature was 150°C and drying gas flow rate was 13 L/min. 5 μL of sample was injected onto an Agilent Zorbax RRHD Extend-C18 (1.8 μm, 2.1 × 150 mm) column maintained at 35°C. A 24 min chromatographic gradient was performed using mobile phase A and mobile phase B (10 mM tributylamine and 15 mM acetic acid in methanol) at a flow rate of 0.25 mL/min. Following the 24 min gradient the analytical column was backflushed for 6 min with 99% acetonitrile at a 0.8 mL/min flow rate followed by a 5 min re-equilibration step (MassHunter Metabolomics dMRM Database and Method, Agilent Technologies). Data analysis was performed with MassHunter Quantitative Analysis (v. B.09.00). Peak areas for each metabolite were normalized to d5-2HG and to protein content of matched duplicate samples as determined by bicinchoninic acid (BCA) assay (Thermo Fisher Scientific, 23225).

Principal component analysis (PCA) was performed by feeding a matrix of normalized metabolite peak areas (as described above) to the prcomp function in R (v. 4.3.2) with the “scale” argument set to “TRUE”. Partial least squares-discriminant analysis (PLS-DA) and variable importance in the projection (VIP) analysis were performed using MetaboAnalyst 6.0^63^. The Venn diagram showing overlapping VIP results was generated using the R package eulerr (version 7.0.2). For principal component analysis integrating separate experiments, z-scores were first calculated within each experiment using the scale function in R (v. 4.3.2) with default parameters. PCA was then performed with the prcomp function in R with default parameters.

### Western Blotting

Protein lysates were extracted in RIPA buffer (Cell Signaling Technology, 9806) and concentration determined by BCA assay (Thermo Fisher Scientific, 23225). Samples were boiled for 5 min, then separated by SDS-polyacrylamide gel electrophoresis and transferred to nitrocellulose membranes (Bio-Rad, 1620115). Membranes were blocked in 3% milk in Tris-buffered saline with 0.1% Tween-20 (TBST) followed by incubation with primary antibodies at 4°C overnight. Following TBST washes, membranes were incubated with horseradish-peroxidase-conjugated secondary antibodies (Cytiva, mouse, NA931; rabbit, NA934) for at least 1 h at room temperature, incubated with Pierce ECL Western Blotting Substrate (Thermo Fisher Scientific, 32109) and imaged using HyBlot CL Autoradiography Film (Denville Scientific, E3018) with an SRX-101A X-ray Film Processor (Konica Minolta). Antibodies used were anti-DHFR (1:1000, Cell Signaling Technology, 45712), anti-TYMS (1:1000, Cell Signaling Technology, 9045), anti-FOXO1 (1:1000, Cell Signaling Technology, 2880), and anti-vinculin (1:10,000, Sigma, V9131).

### EdU incorporation flow cytometry assay

Cells were seeded in triplicate into 12-well plates. 24 h after seeding, cells were changed to treatment media containing DMSO (Sigma, D2650) or 0.2 μM methotrexate (Sigma, M9929). Cells were collected for flow cytometry analysis 48 h after starting treatment. One hour before collection, cells were changed into fresh medium containing 10 μM EdU (Vector Laboratories, CCT-1149). Cells were collected and stained with ZombieGreen Fixable Viability dye (Biolegend, 423111), followed by fixation with 4% paraformaldehyde and permeabilization with saponin-based permeabilization and wash reagent (Thermo Fisher Scientific, C10419). Fixed and permeabilized cells were stained using the Click-iT Cell Reaction Buffer Kit (Thermo Fisher Scientific, C10269) and AZDye 405 Picoyl Azide (Vector Laboratories, CCT-1308) according to the manufacturer’s instructions. Samples were analyzed on an LSRFortessa flow cytometer using FACSDiva v8.0 (BD Biosciences). Analysis of EdU incorporation was performed with FlowJo (version 10.10.0).

### RNA-sequencing and RNA-sequencing analysis

RNA was isolated with TRIzol (Thermo Fisher Scientific, 15596018) according to manufacturer’s instructions. After RiboGreen quantification and quality control by Agilent BioAnalyzer, 500 ng of total RNA with RIN values of 9.3-10 underwent polyA selection and TruSeq library preparation according to instructions provided by Illumina (TruSeq Stranded mRNA LT Kit, catalog #RS-122-2102), with 8 cycles of PCR. Samples were barcoded and run on a NovaSeq 6000. FASTQ files were aligned to the hg38 reference genome using Dragen v3.10 (Illumina) to generate BAM files. A matrix of raw counts was generated using featureCounts/subread (version 2.16.1)^64^. Differentially expressed genes were determined using DESeq2 (version 1.42.1)^65^. Overrepresentation analysis was performed using clusterProfiler (version 4.10.1)^66^. To assess correlation of gene expression between MTX data and KFR shPAX3-FOXO1 data (obtained from GSE218974, see below), the Pearson correlation coefficient and corresponding p-value were calculated using the cor.test function in R (version 4.3.2).

### Published RNA-sequencing analysis

For the published RH30 PAX3-FOXO1 degron data^24^, FASTQ files were downloaded from the Gene Expression Omnibus (GSE183281) using SRAToolkit (version 3.1.0) and filtered using fastp (version 0.23.4)^67^. Filtered FASTQ files were aligned to the hg38 reference genome using STAR (version 2.7.11b)^68^ and a matrix of raw counts was generated using featureCounts/subread (version 2.16.1)^64^. Differentially expressed genes were determined using DESeq2 (version 1.42.1)^65^. Hierarchical clustering was performed using code adapted from a publicly available repository on Github (https://tavareshugo.github.io/data-carpentry-rnaseq/04b_rnaseq_clustering.html). To select candidate genes for hierarchical clustering, a significance cutoff of *P_adj_* < 0.05 was used.

For published RMS patient tumor data^50,69^, a raw counts matrix was downloaded from the Gene Expression Omnibus (GSE108022). Gene signature scores were calculated by adding the z-scores for all genes in the stated list and dividing by the square root of the gene list size^70^.

### DepMap mRNA analysis

Gene expression data were downloaded from the DepMap portal, 24Q2 release^22^. To compare gene expression between fusion-negative (*n* = 10) and PAX3-FOXO1^+^ (*n* = 7) RMS cell lines, average expression values (using reported transcript per million (TPM) values) were calculated for every gene within each subtype, and the fusion-negative averages were subtracted from the PAX3-FOXO1^+^ averages. Genes were ranked in descending order of difference, and this ranked list was used as input for GSEA.

### DepMap differential essentiality analysis

Gene essentiality data were downloaded from the DepMap portal, 24Q2 release^22^. To calculate differential essentiality between fusion-negative (*n* = 5) and PAX3-FOXO1^+^ (*n* = 7) RMS cell lines, the average essentiality score for every gene was calculated within each subtype, the PAX3-FOXO1^+^ averages were subtracted from the fusion-negative averages for each gene, and genes were ranked in descending order of difference.

### Single cell RNA-sequencing analysis

Published data^20^ for KFR cells expressing a doxycycline-inducible hairpin against PAX3-FOXO1 were downloaded from the Gene Expression Omnibus (GSE218974). Data were processed with Seurat (version 5.1.0)^71^ following the standard pipeline. To generate gene signature scores, the AddModuleScore function was utilized with the indicated gene lists in Supplementary Table 3.

### Statistics and reproducibility

All statistical analyses were performed in Prism 10 (GraphPad) except for DepMap differential essentiality analysis, RNA-seq overrepresentation analysis, and hierarchical clustering, which were performed in R (v4.3.2), and GSEA, which was performed using GSEA software (v4.3.3). No statistical methods were used to pre-determine sample size. All *in vitro* experiments were independently repeated at least twice with similar results.

### Data and reagent availability

RNA-sequencing data generated by this study have been deposited in the Gene Expression Omnibus under the accession code GSE286950. Previously published bulk and single-cell RNA-sequencing data utilized or reanalyzed for this study are available under the accession codes GSE183281, GSE108022, and GSE218974. DepMap data (24Q2 release) were downloaded from the DepMap Portal (https://depmap.org/portal). All unique reagents generated by this study will be made available, following request, from the corresponding author with a completed material transfer agreement.

## Acknowledgements

We thank members of the Finley lab, Santosh Vardhana, Jovana Pavisic, and Damon Reed for discussion and Andrew Intlekofer for use of shared LC-MS instrumentation. K.I.P. was supported by a Bruce Charles Forbes Pre-Doctoral Fellowship (MSKCC). J.S.B. was supported by a Human Frontier Science Program Fellowship (L1000200/2021-L) and is a Kravis WiSE Fellow (MSKCC). A.M.M. is supported by a T32 training grant from the NICHD (T32HD060600-15). B.T.J. is supported by a Ruth L. Kirschstein Predoctoral fellowship from the NICHD (no. F30HD107943). S.C. is an MSK Bridge Scholar and Damon Runyon SPARK Scholar. A.X. is supported by a Ruth L. Kirschstein Predoctoral fellowship from the NCI (no. F30CA284711-01). B.T.J. and A.X. were additionally supported by a Medical Scientist Training Program grant from the NIGMS of the National Institutes of Health under award number T32GM152349 to the Weill Cornell/Rockefeller/Sloan Kettering Tri-Institutional MD-PhD Program. L.W.S.F. is a New York Stem Cell Foundation – Robertson Investigator. We acknowledge the use of the Integrated Genomics Operation Core, funded by the NCI Cancer Center Support Grant (CCSG, P30 CA08748), Cycle for Survival, and the Marie-Josée and Henry R. Kravis Center for Molecular Oncology. This work was additionally supported by grants from the NYSCF, the Pershing Square Sohn Foundation, the Geoffrey Beene Cancer Research Center (to L.W.S.F.), the PaulieStrong Foundation (to F.D.C.), and the Memorial Sloan Kettering Cancer Center Support Grant (P30CA08748).

## Authors’ Contributions

K.I.P. and L.W.S.F. conceived the study. K.I.P. carried out experiments with assistance from J.S.B., A.M.M., A.X., S.C., and L.P. K.I.P. conducted all computational analysis with assistance from B.T.J. K.G., A.S., D.Y., and F.D.C. assisted with PDX implantation and study design. J.A.B. and J.D.R. assisted with nucleotide analyses. K.I.P. and L.W.S.F. wrote the manuscript with input from all authors.

---

**Supplementary Figure 1.**
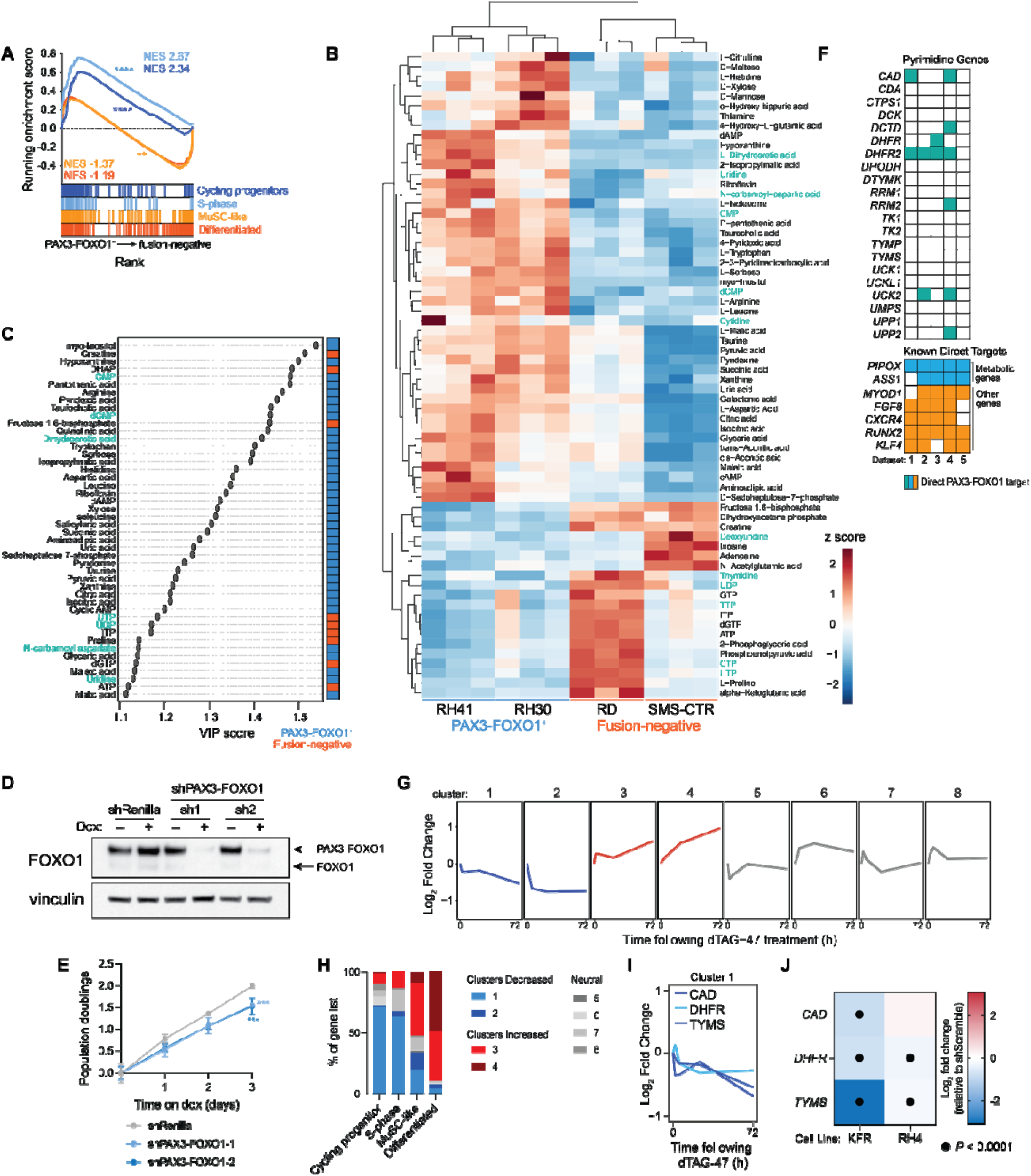
**A**, Gene set enrichment analysis (GSEA) of published RNA-sequencing data (DepMap 24Q2)^22^ comparing expression of cell fate gene lists in PAX3-FOXO1^+^ (*n* = 7) and fusion-negative RMS (*n* = 10) cell lines. **B**, Heatmap of all 66 metabolites with significantly (*P* < 0.05) different levels between PAX3-FOXO1^+^ and fusion-negative RMS cell lines at baseline. Pyrimidine metabolites are highlighted in teal. **C**, Variable importance in the projection (VIP) analysis of metabolites discriminating between PAX3-FOXO1^+^ and fusion-negative RMS cell lines at baseline. Pyrimidine metabolites are shown in teal. Column to the right indicates whether a metabolite’s levels are higher in PAX3-FOXO1^+^ RMS cell lines (blue) or fusion-negative RMS cell lines (orange). Data are from same experiment as data shown in **B**. **D**, Western blot of PAX3-FOXO1 (top band detected by FOXO1 antibody) in RH30 cells expressing shRNAs against *Renilla* luciferase (control) or *PAX3-FOXO1.* Cells were cultured with doxycycline (Dox) for 72 h to induce shRNA expression. **E**, Population doublings of RH30 cells expressing shRenilla (control) or shPAX3-FOXO1, cultured with doxycycline (Dox) to induce shRNA expression over 72 h. **F**, Table showing whether pyrimidine genes are consistent direct targets of PAX3-FOXO1, compared to known direct targets. Filled-in boxes indicate that the gene was identified as a direct target of PAX3-FOXO1 in that dataset. Data were obtained from published ChIP-sequencing of endogenous PAX3-FOXO1 in RH30 (dataset 1)^25^ or RH4 (datasets 2-3)^25,26^, or PAX3-FOXO1 overexpression in Dbt immortalized myoblasts (dataset 4)^27^ or CRL7250 fibroblasts (dataset 5)^27^. **G**, Eight gene clusters that result from hierarchical clustering analysis of published RNA-sequencing of RH30-PAX3-FOXO1 degron system^24^. **H**, Bar graph showing distribution of each gene set across the clusters in **G**. Analysis includes genes that met inclusion criteria (expression significantly (*P* < 0.05) changed by PAX3-FOXO1 degradation for at least one timepoint) for hierarchical clustering. **I**, Expression of *DHFR*, *TYMS*, and *CAD* upon PAX3-FOXO1 degradation. All three genes are found in Cluster 1 from **G**. **J**, Heatmap depicting published expression data of *DHFR, TYMS,* and *CAD* in two PAX3-FOXO1^+^ cell lines (KFR, RH4) expressing shRNA against *PAX3-FOXO1*^20^. Data are displayed as log_2_ fold change relative to shScramble (control) and represent averaged expression values of all cells. For **E**, significance was assessed by 2-way ANOVA with Dunnett’s multiple comparisons test comparing shRenilla (control) to each shPAX3-FOXO1 line. Significance displayed for the 72 h timepoint. (\*\*\**P* < 0.001). Gene lists are provided in Supplementary Table 3.

**Supplementary Figure 2.**
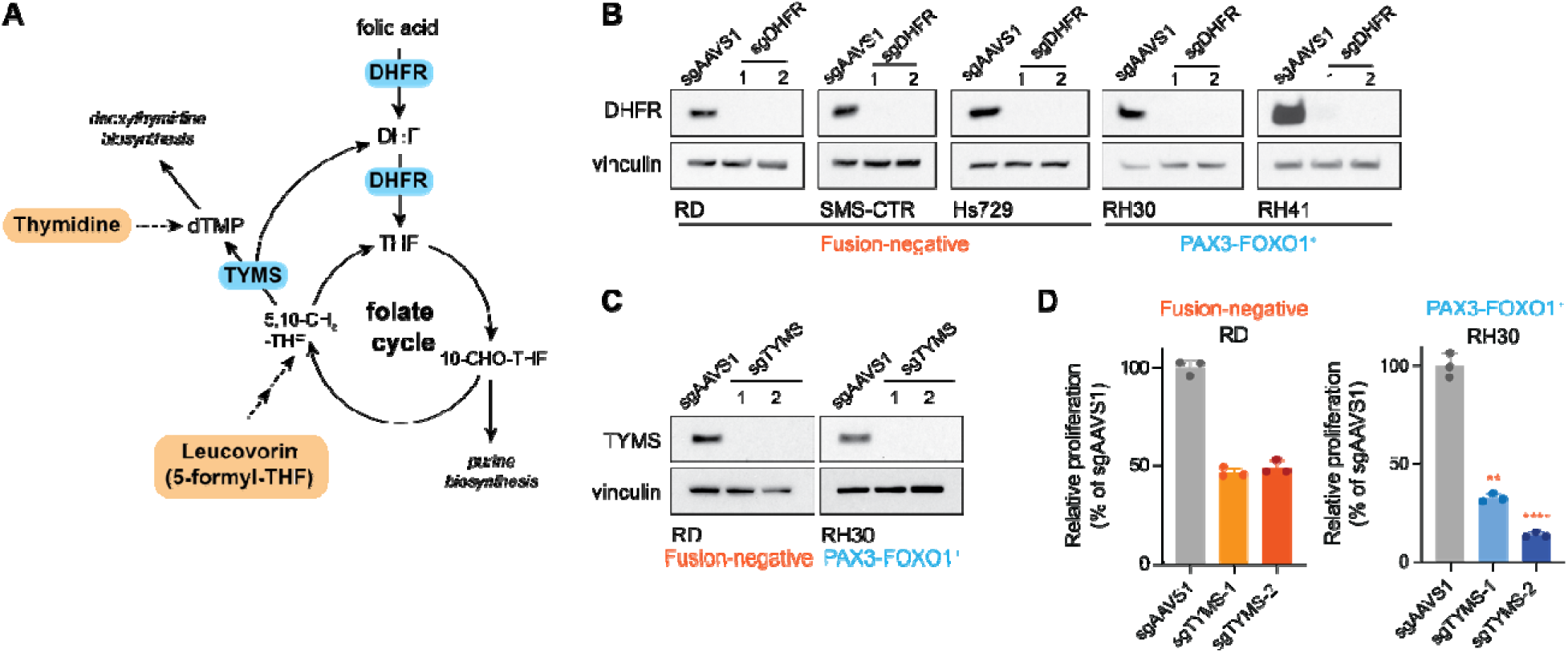
**A**, Schematic of folate cycle depicting leucovorin (rescue agent for loss of DHFR) and thymidine (rescue agent for loss of TYMS, along with cytidine). **B**, Western blots of RMS cells with control (sgAAVS1) or *DHFR*-editing (sgDHFR-1 and sgDHFR-2). **C**, Western blots of RMS cells with control (sgAAVS1) or *TYMS*-editing (sgTYMS-1 and sgTYMS-2). **D**, Relative proliferation of RMS cells in response to *TYMS* editing using two independent sgRNAs (sgTYMS-1, sgTYMS-2). Data are shown as a percentage of control (sgAAVS1) for each cell line, and are mean ± s.d., *n* = 3 independent replicates. Significance was assessed using one-way ANOVA with Tukey’s multiple comparisons test, and for each sgRNA, RH30 (PAX3-FOXO1^+^) was compared to RD (fusion-negative). (\*\**P* < 0.01, \*\*\*\**P* < 0.0001)

**Supplementary Figure 3.**
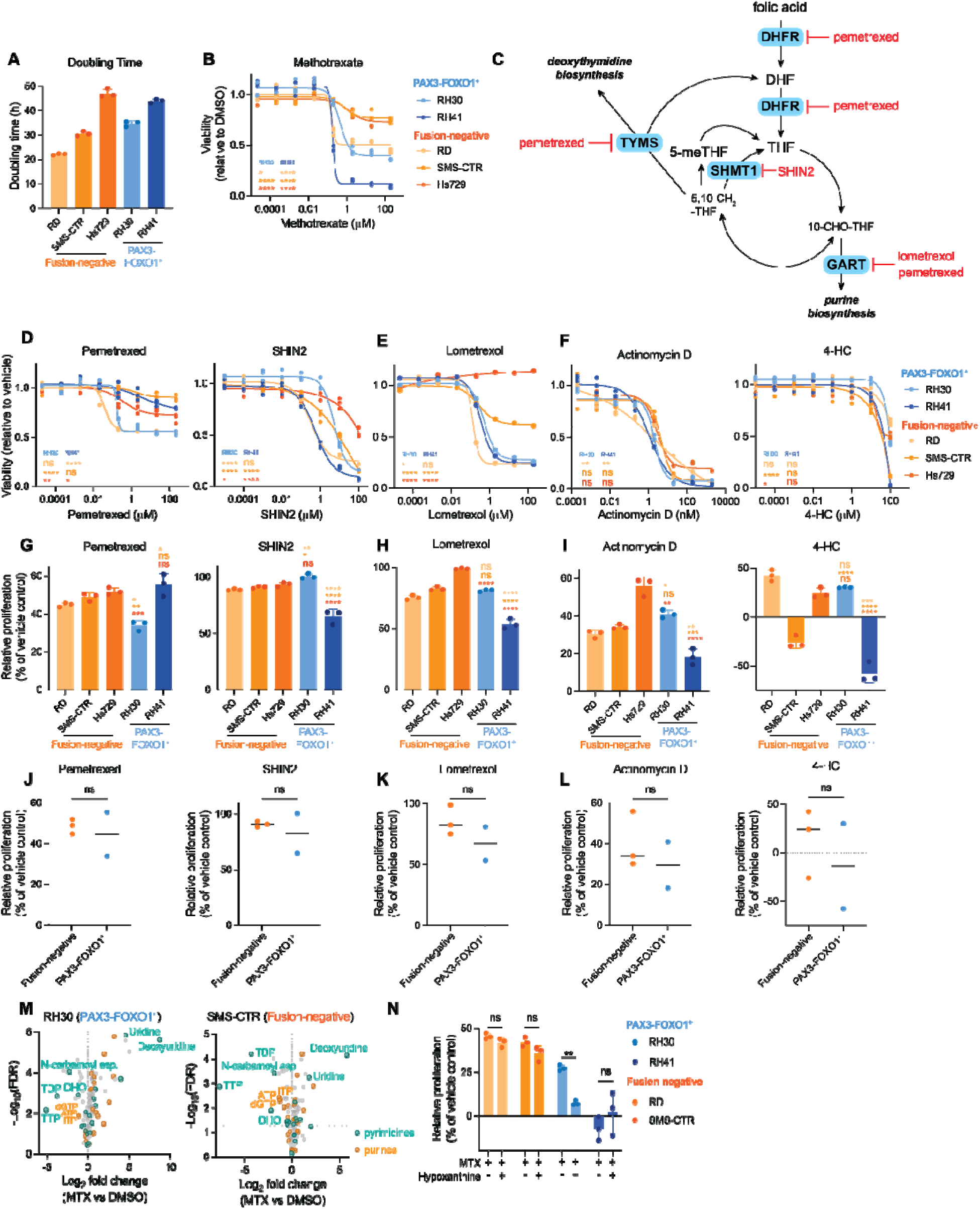
**A**, Doubling times (in h) for RMS cell lines, calculated over 96 h. **B**, Dose response curve for MTX in RMS cell lines cultured in medium with 5-meTHF as the folate source measured by CellTiter-Glo viability assay after 48 h of treatment. Maximum dose tested was 200 µM. **C**, Schematic depicting the cytosolic folate cycle with targets of pemetrexed, SHIN2, and lometrexol. (SHMT1 = serine hydroxymethyltransferase 1, GART = phosphoribosylglycinamide formyltransferase) **D-F**, Dose response curves for pemetrexed (**D**), SHIN2 (**D**), lometrexol (**E**), actinomycin D (**F**), and 4-HC (**F**) in RMS cell lines, measured by CellTiter-Glo viability assay after 48 h of treatment. Maximum doses tested were 200 µM (pemetrexed, lometrexol), 100 µM (SHIN2, 4-HC), or 2 µM (actinomycin D). **G-I**, Relative proliferation of RMS cells in response to pemetrexed (**G**, 2 µM), SHIN2 (**G**, 2 µM), lometrexol (**H**, 2 µM), actinomycin D (**I**, 0.5 nM), or 4-HC (**I**, 20 µM) for 96 h. Data are shown as a percentage of the vehicle control (DMSO for all except pemetrexed, which is H_2_O). **J-L**, Summary of the effects of each drug on relative proliferation shown in **G-I**. Each dot represents the average of all replicates from each cell line. **M**, Volcano plots for two RMS cell lines (RH30, PAX3-FOXO1^+^ and SMS-CTR, fusion-negative) showing the log_2_ fold change in metabolite abundance in cells treated with MTX (0.2 µM, 48 h) compared to cells treated with vehicle (DMSO). Dotted horizontal line indicates significance threshold FDR < 0.05. **N**, Relative proliferation of RMS cells in response to MTX (0.2 µM) in the presence or absence of hypoxanthine (100 µM) for 96 h. Data are shown as a percentage of the vehicle control (DMSO and 0.1N NaOH). For **A**, **B**, **D**-**I**, and **N**, data are mean ± s.d., *n* = 3 independent replicates. Significance was assessed by ordinary one-way ANOVA with Tukey’s multiple comparisons test (**B**, **D**-I), 2-way ANOVA with Šídák’s multiple comparisons test (**N**), and unpaired two-tailed Student’s *t*-test (**J**-**L**). In **B** and **D**-**I**, each PAX3-FOXO1^+^ cell line was compared to each fusion-negative cell line. For dose response curves, comparison shown for 0.2 µM (MTX, **B**), 2 µM (pemetrexed, **D** and lometrexol, **E**), 1 µM (SHIN2, **D**), 0.2 nM (actinomycin D, **F**), and 10 µM (4-HC, **F**). For **N**, treatment groups (MTX or MTX + hypoxanthine) were compared to each other within each cell line. (ns = not significant, \**P* < 0.05, \*\**P* < 0.01, \*\*\**P* < 0.001, \*\*\*\**P* < 0.0001)

**Supplementary Figure 4.**
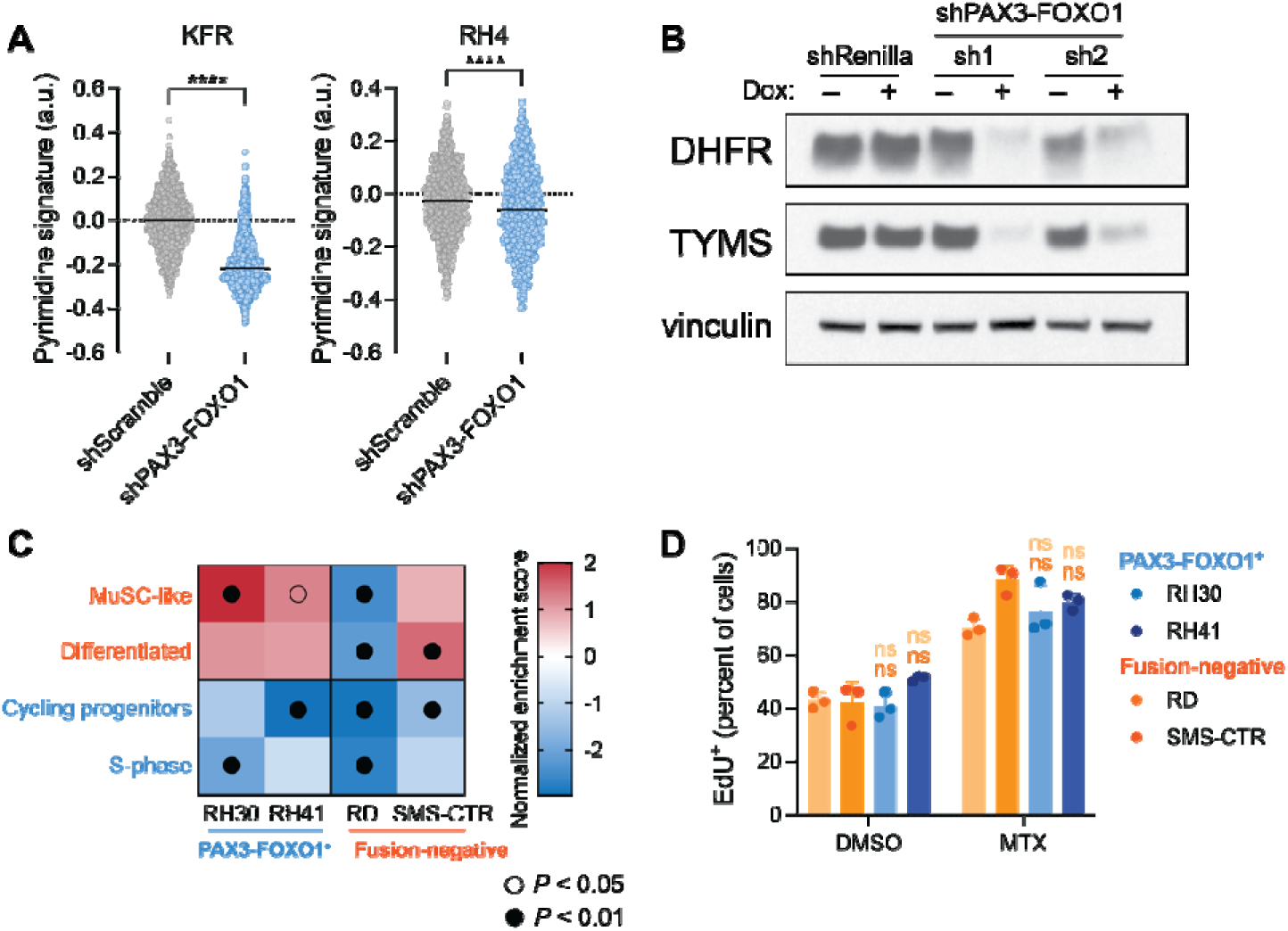
**A**, Violin plots of pyrimidine gene signature scores in two independent PAX3-FOXO1^+^ cell lines (KFR, RH4), obtained from published single-cell RNA-sequencing data^20^. Cells were engineered to express shScramble (control) or shPAX3-FOXO1. Lines represent the median of each group. **B**, Western blot of DHFR and TYMS in RH30 cells expressing shRNAs against *Renilla* luciferase (control) or *PAX3-FOXO1.* Cells were cultured with doxycycline (Dox) for 72 h to induce shRNA expression. **C**, Heatmap of normalized enrichment scores generated by GSEA performed on four RMS cell lines treated with MTX (0.2 µM) for 48 h. Gene lists are provided in Supplementary Table 3. **D**, Percentage of cells with EdU incorporation after treatment with vehicle (DMSO) or MTX (0.2 µM) for 48 h, measured via flow cytometry. Data are mean ± s.d., *n* = 3 independent replicates. Significance was assessed by unpaired two-tailed Student’s *t* test comparing shScramble and shPAX3-FOXO1 (**A**) or ordinary one-way ANOVA with Tukey’s multiple comparisons test (**D**), with each PAX3-FOXO1^+^ cell line compared to each fusion-negative cell line in DMSO and MTX conditions. (ns = not significant, \*\*\*\**P* < 0.0001)

**Supplementary Figure 5.**
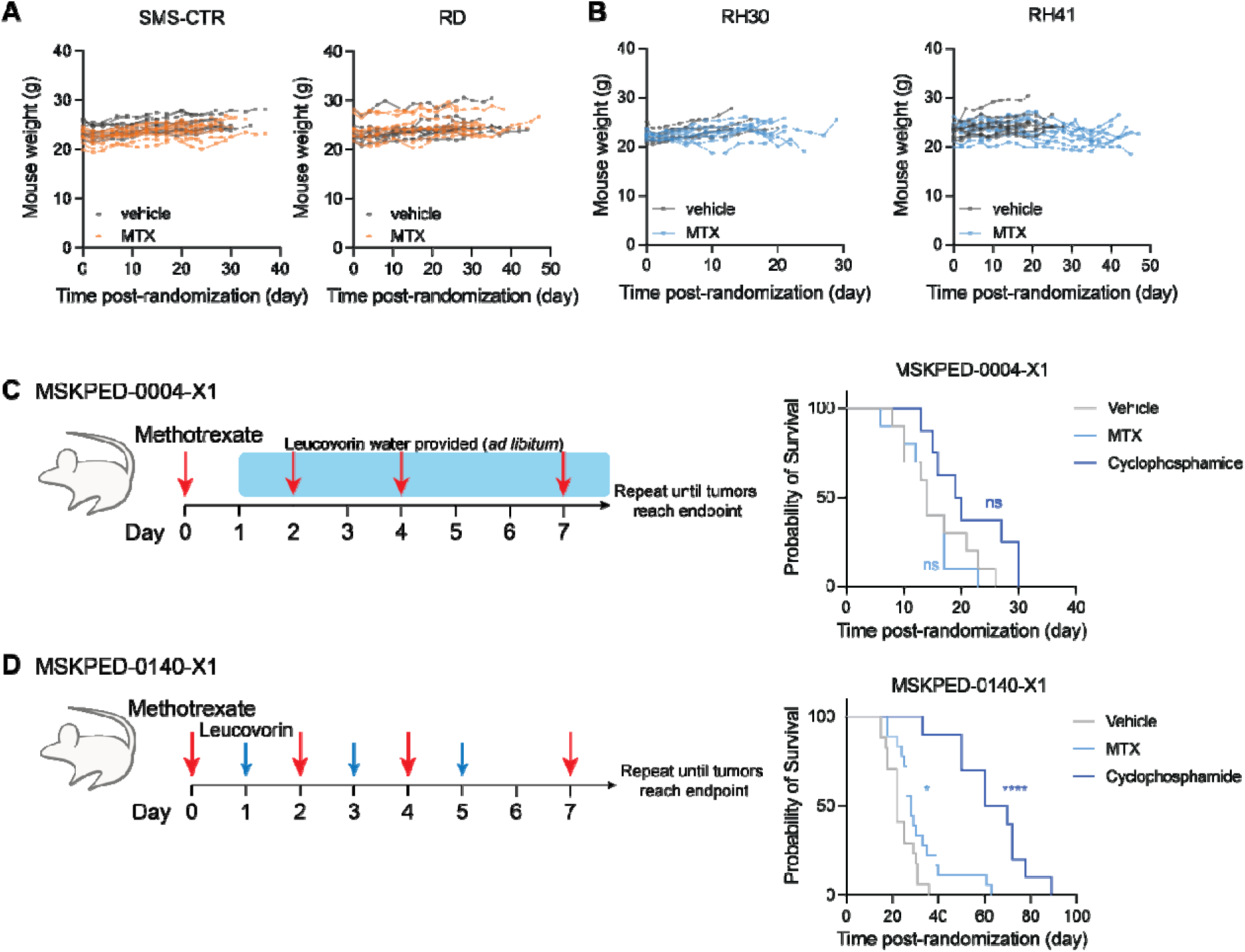
**A**, **B**, Weights of mice bearing fusion-negative RMS (**A**, SMS-CTR and RD) or PAX3-FOXO1^+^ RMS (**B**, RH30 and RH41) cell line xenografts during treatment with vehicle (corn oil) or MTX. **C**, (Left) Schematic depicting MTX and leucovorin dosing strategy for MSKPED-0004-X1, in which leucovorin was continuously provided to mice starting 24 h after the first dose of MTX was administered. (Right) Survival curve for mice bearing PAX3-FOXO1^+^ PDX MSKPED-0004-X1 and treated with vehicle (corn oil), MTX, or cyclophosphamide. Significance was assessed by log-rank (Mantel-Cox) test. **D**, (Left) Schematic depicting MTX and leucovorin dosing strategy for MSKPED-0140-X1, in which leucovorin was administered 24 h after each dose of MTX. (Right) Survival curve for mice bearing PAX3-FOXO1^+^ PDX MSKPED-0140-X1and treated with vehicle (corn oil), MTX, or cyclophosphamide. Significance was assessed by log-rank (Mantel-Cox) test. For MSKPED-0004-X1 (**C**), *n* = 8 (vehicle, cyclophosphamide) or 9 (MTX). For MSKPED-0140-X1 (**D**), *n* = 17 (vehicle, MTX) or 11 (cyclophosphamide). (ns = not significant, \**P* < 0.05, \*\*\*\**P* < 0.0001)

## Notes

### Competing Interest Statement

The authors have declared no competing interest.

